# A cellular automaton model of osteogenic differentiation reveals identifiability limits of endpoint assays

**DOI:** 10.64898/2026.04.23.720356

**Authors:** Ali Aslan Demir, Thomas Combriat, Catherine Anne Heyward, Hanna Tiainen, Aurélie Carlier, Dag Kristian Dysthe

## Abstract

Standard differentiation assays sample cell states only at discrete time points, while the underlying progression unfolds continuously and heterogeneously across cells. As a result, different combinations of proliferation, commitment, and maturation dynamics can converge to similar endpoint measurements. This many-to-one mapping between latent trajectories and observable readouts constitutes a partially observed inverse problem that limits mechanistic interpretation. Although this ambiguity is inherent to many experimental systems, it is rarely examined using models that connect cell-state dynamics to assay-level quantities.

We present OsteoMin, a coarse-grained cellular automaton that links stochastic transitions between pre-osteoblast and osteoblast states to experimentally measurable readouts of alkaline phosphatase activity, collagen deposition, and mineralization. Model parameters were constrained using literature-reported kinetics and evaluated against dexamethasone and menaquinone-4 perturbations. The framework reproduces qualitative assay trends and enables systematic analysis of how cell-state progression, matrix maturation, and external perturbations shape differentiation outcomes.

Using this framework, we quantify the identifiability limits of endpoint assays and test whether standard measurements can distinguish underlying differentiation mechanisms. Distinct perturbation families often produce similar endpoint responses (macro-F1 ≈ 0.42), indicating limited discriminative power. Incorporating temporal trajectories improves separability (macro-F1 ≈ 0.78), demonstrating that most identifiable information resides in marker dynamics rather than terminal measurements. Sobol analysis shows early markers depend on proliferation timing, whereas late mineralization is governed by nonlinear matrix maturation and parameter interactions. Together, these results show that endpoint assays constrain overall progression but do not uniquely identify underlying mechanisms. OsteoMin provides a framework linking differentiation dynamics to assay observables and a basis for assessing identifiability in endpoint-driven systems.

## 1 Introduction

Biomineralization describes the formation and organization of inorganic phases in biological systems [1]. In mammalian bone, this process is driven by osteogenic differentiation, a regulated program that has been studied extensively for decades [2, 3]. MC3T3-E1 mouse calvarial pre-osteoblasts are widely used as an in vitro model of osteogenesis because they mineralize consistently under defined culture conditions [4]. Because MC3T3-E1 cells are lineage committed, variation in initial cell-state composition can alter differentiation trajectories and contribute to experimental variability [5].

Experimental characterization of osteogenic differentiation typically relies on endpoint assays that probe early, intermediate, and late stages of the program, including alkaline phosphatase (ALP) activity, collagen type I deposition, and mineralization measured by Alizarin Red S staining (ARS) [2, 6–9]. These measurements are population-averaged and collected at discrete time points, and many assays are destructive, preventing repeated observation of the same cells over time [10]. As a result, experimental data provide sparse and indirect observations of an underlying continuous differentiation process, limiting access to its dynamics and multi-scale interactions [11]. In parallel, lineage commitment, matrix deposition, and mineral nucleation proceed asynchronously within MC3T3-E1 cultures [12–14].

Understanding differentiation dynamics requires linking cell-level processes such as proliferation, migration, and state transitions to experimentally observable outcomes. Cell-based computational models provide a framework for connecting individual behaviors to population-level readouts [15]. Cellular automaton (CA) models represent multicellular systems using rule-based descriptions of cell processes and have been applied to growth and mineralization-related phenomena [16–19]. Existing in silico osteogenesis models typically reproduce temporal marker trajectories or spatial mineral patterns but do not explicitly examine whether standard assay panels contain sufficient information to distinguish alternative mechanistic hypotheses of lineage progression [20]. Because these readouts provide only partial observations of the underlying process, the relationship between observable markers and differentiation dynamics constitutes a partially observed inverse problem [21]. In such systems, multiple parameter combinations can yield similar observable behavior, leading to a many-to-one mapping between underlying dynamics and measured outputs [22].

Together, sparse endpoint sampling and many-to-one model behavior raise a central question: to what extent can standard endpoint measurements resolve underlying differentiation mechanisms? We frame this as a problem of partial observability in a stochastic cell-state system.

We study osteogenic differentiation under controlled media conditions. Standard osteogenic medium contains ascorbic acid (AA), which supports collagen synthesis, and *β*-glycerophosphate (*β*-GP), which provides phosphate for mineralization [23]. We introduce controlled perturbations using dexamethasone (Dex) and menaquinone-4 (MK-4), which act at partially distinct stages of differentiation: Dex promotes early lineage commitment [24, 25], whereas MK-4 enhances osteogenic gene expression and matrix mineralization at later stages via *γ*-carboxylation–dependent pathways [26, 27]. These stage-specific effects provide a basis to test whether standard assay panels can distinguish mechanistically distinct responses [28].

We develop OsteoMin, a cellular automaton model linking latent MC3T3-E1 differentiation dynamics to assay readouts of ALP activity, collagen deposition, and mineralization. The model connects stochastic cell-state transitions, matrix accumulation, and mineral formation to experimentally accessible observables, enabling direct comparison with experimental measurements.

Using this framework, we examine how cell-state progression, spatial organization, matrix maturation, and media perturbations shape assay outcomes. Literature-parameterized simulations are compared with experimental endpoint data and complemented by module ablation and sensitivity analysis to identify mechanisms governing early and late readouts.

Finally, we assess practical identifiability under endpoint sampling by comparing end-point features with time-resolved trajectories, testing whether standard assay panels can distinguish alternative perturbation families and quantifying how strongly measurements constrain underlying dynamics.

## 2 OsteoMin Cellular Automaton Model

### 2.1 Overview of the OsteoMin model

OsteoMin was designed to establish a structural and interpretable link between latent osteogenic cell-state progression and commonly used endpoint assays. Rather than reproducing differentiation only at the level of generic growth dynamics or mineral pattern formation, the model directly connects stochastic transitions between pre-osteoblast and osteoblast states to experimentally measurable readouts. By aligning simulated outputs with assay observables, the framework enables mechanistic interpretation of endpoint measurements while also allowing assessment of how strongly such measurements constrain the underlying differentiation dynamics.

OsteoMin represents MC3T3-E1 monolayer cultures as a two-dimensional cellular automaton in which lattice sites are either empty or occupied by a single agent corresponding to a pre-osteoblast (pre-OB) or osteoblast (OB). Cells follow rule-based behaviors of migration, proliferation, differentiation, and apoptosis, while producing local extracellular fields corresponding to ALP activity, collagen matrix, and mineral (hydroxyapatite; HAp). These marker fields accumulate over time and are aggregated across the lattice to produce simulated assay readouts comparable to experimental ALP activity measurements, collagen immunofluorescence, and Alizarin Red S staining. The update scheme is illustrated in Fig. 1.

**Fig 1:**
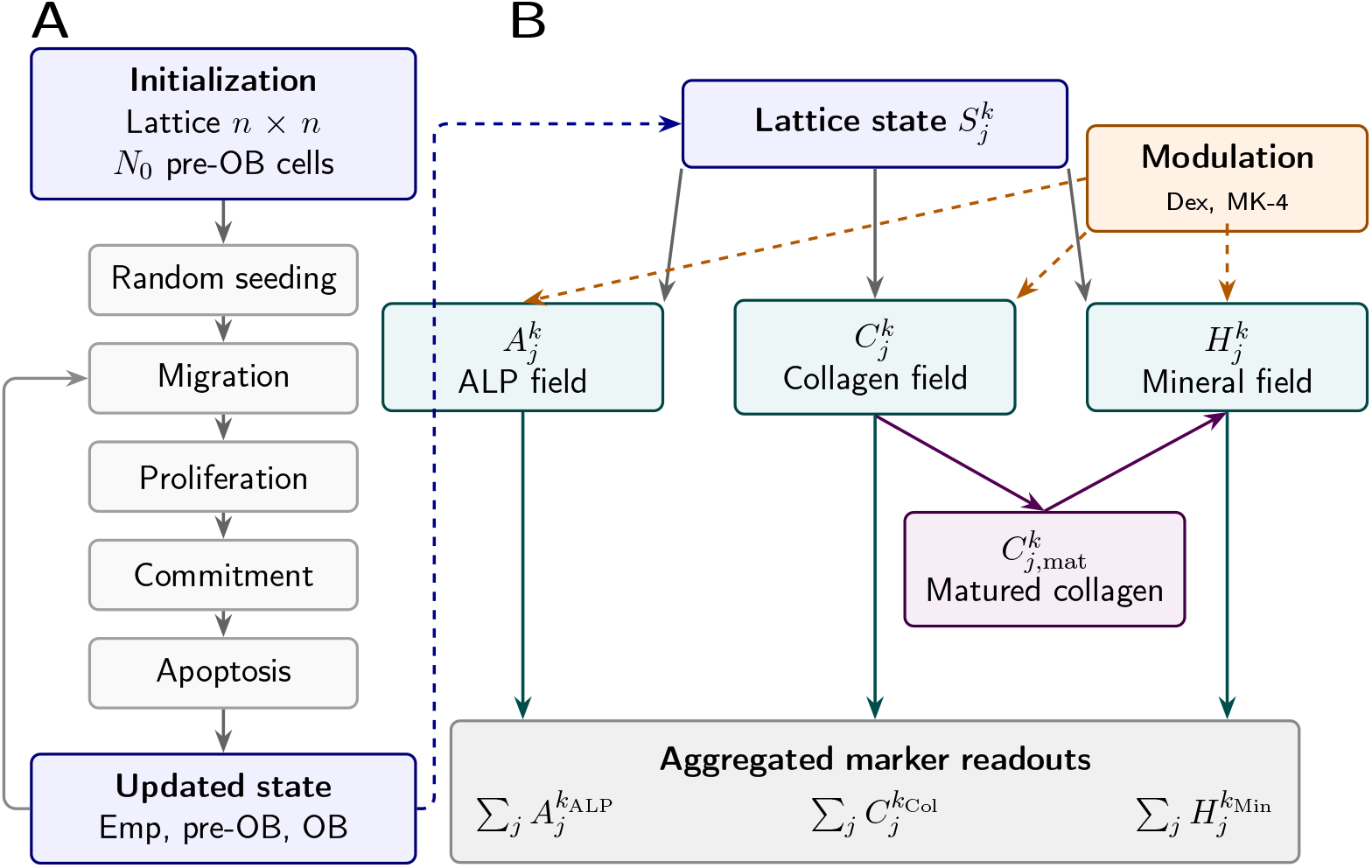
Schematic of the OsteoMin model. (**A**) Cellular automaton update sequence for pre-OB and OB states, including migration, proliferation, commitment, and apoptosis. (**B**) The lattice state generates local ALP, collagen, and mineral fields, with Dex and MK-4 modulating osteogenic production. Aggregation of these fields yields simulated assay readouts comparable to experimental ALP, collagen, and mineralization endpoints.

### 2.2 Spatial and Temporal Representation

The simulation domain is a square lattice of size *n* × *n*, with a default grid of 33 × 33 sites corresponding to an area of approximately 1 × 1 mm. Each lattice site represents the footprint of a single cell with spacing Δ*x* ≈ 30 µm, consistent with the diameter of MC3T3-E1 cells. Periodic boundary conditions are applied. Time is discretized as *t*_*k*_ = *k*Δ*t* with timestep Δ*t* = 1 h, chosen to match reported migration timescales.

State updates follow a sequential asynchronous scheme. Cellular updates are applied in fixed order (movement, proliferation, commitment, apoptosis, marker production), while cells within each class are visited individually in randomized order at each timestep to minimize ordering bias.

Updates take effect immediately within each stage. During proliferation, target sites are reserved immediately, while daughter cells become active in the subsequent timestep. Cells undergoing apoptosis are removed before marker-field updates, ensuring extracellular quantities reflect the updated cell population.

Stochastic events were generated using NumPy’s pseudo-random number generator. Simulations used a fixed initial seed (12345) unless otherwise stated. For experimental– simulation comparisons (Section 4.5) and latent initialization analyses (Section 4.3), in-dependent randomly initialized seeds were used (*n* = 10 per condition).

### 2.3 Cell State Dynamics

Pre-OBs are seeded at *t*_0_ = 0 and can move, proliferate, and commit to the OB lineage. OBs remain stationary and deposit increased levels of assay markers. Apoptotic cells are removed from the lattice, leaving empty sites that can be occupied by pre-OBs.

#### 2.3.1 Initialization

Simulations start by placing *N*_0_ pre-OB cells on the lattice using random placement.

#### 2.3.2 Migration

Pre-OBs perform a stochastic random walk on a square lattice (Δ*t* = 1 h), attempting a move to a randomly selected empty Moore-neighbor site with probability *p*_*m*_. Axial and diagonal directions are weighted to reduce lattice anisotropy and improve the rotational symmetry of long-time diffusion.

The migration probability *p*_*m*_ was calibrated using migration data for MC3T3-E1 cells on flat substrates reported by Refaaq et al. [29]. The mean-squared displacement increases linearly with time, yielding an effective diffusion coefficient of ~ 300 − −320 µm^2^ h^−1^. For a lattice spacing Δ*x* = 30 µm, matching this hourly rate led to *p*_*m*_ ≈ 0.27. Derivation of the migration mapping is provided in S1 Text, Section 1.2.

#### 2.3.3 Proliferation

Each pre-OB is assigned an individual division period *τ*_*i*_ ∈ [38, 52] h at birth [30]. To represent cell-cycle progression, each cell carries an internal age variable 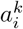 measuring the time since its last division. Cell age is updated at each timestep as 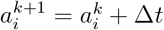.

When 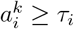, the cell attempts to divide by placing a daughter pre-OB in a randomly selected empty Moore-neighbor site. Upon successful division, the parent cell age is reset to zero and the daughter cell is assigned a new division period *τ*_*i*_ drawn independently. If no neighboring site is available, division is postponed and retried in subsequent timesteps.

#### 2.3.4 Osteogenic commitment

Commitment from pre-OB to OB was modeled as a delayed stochastic transition. Following osteogenic induction at time *t*_osteo_, cells remain uncommitted for a maturation delay *t*_*d*_. After this delay (*t*_*k*_ ≥ *t*_osteo_ + *t*_*d*_), each eligible pre-OB transitions irreversibly to the OB state with probability *p*_commit_ = 1 − exp(−*k*_commit_ Δ*t*) per timestep, where *k*_commit_ is the commitment rate.

The parameters *t*_*d*_ and *k*_commit_ were chosen to reproduce reported MC3T3-E1 differentiation kinetics, capturing the delay between osteogenic induction, early ALP activity, and subsequent mineralization [3, 31]. Additional implementation details are provided in S1 Text, Section 1.3.

#### 2.3.5 Osteoblast activity

OBs do not proliferate or move. They produce elevated levels of ALP, collagen, and mineral, which accumulate locally at the grid site.

#### 2.3.6 Apoptosis

Apoptosis was modeled as a stochastic death process affecting pre-OBs. At each timestep, pre-OBs undergo apoptosis with probability *p*_*a*_ = 1 × 10^−3^, representing low background turnover in culture [32]. Cells undergoing apoptosis are removed within the same timestep, while previously deposited matrix remains unchanged.

OB apoptosis was not included due to its low reported prevalence (0.6%) and limited quantitative constraints in vitro, which would introduce poorly identifiable parameters [32].

### 2.4 Field dynamics and compound modulation

At each lattice site *j*, three state-associated quantities are tracked: ALP activity 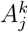, collagen matrix 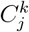, and mineral content 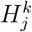 (representing HAp accumulation) at discrete times *t*_*k*_ = *k*Δ*t*. These fields represent cumulative marker production associated with osteogenic differentiation and are aggregated across the lattice to obtain simulated assay readouts. All quantities are expressed in arbitrary units (AU), reflecting relative assay signal magnitudes.

ALP activity is primarily associated with cell-bound matrix vesicles. Marker dynamics are therefore treated as local, and diffusion and degradation are not explicitly modeled [33]. Collagen accumulates locally as an extracellular matrix scaffold that supports mineral nucleation near OB regions [7, 12].

Ascorbic acid (AA) is modeled as a temporally varying medium component due to oxidative degradation (half-life ≈ 8 h) [34], with degradation formulation and effective accumulation provided in S1 Text, Section 1.4. *β*-GP was treated as constant due to its excess concentration (10 mM), regular medium exchange, and rapid diffusion relative to mineralization timescales.

Dexamethasone (Dex) and menaquinone-4 (MK-4) are treated as constant extracellular modulators refreshed during medium exchange. Their effects are represented using phenomenological compound–response functions chosen to reproduce literature-reported dose–response relationships within non-cytotoxic ranges (Dex 0–100 nM; MK-4 0–10 µM), including intermediate optima observed for ALP activity, matrix production, and mineralization. Functional forms capture biphasic behavior while avoiding additional parameters that cannot be uniquely constrained from endpoint measurements [25, 27, 35–37].

#### 2.4.1 Lattice state

Each lattice site *j* has state

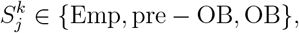

representing empty sites, pre-OBs, and OBs, respectively. Binary occupancy variables indicate live-cell presence and OB identity,

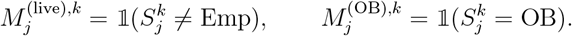

Marker production rates are defined for each marker *i* ∈ {*A, C, H*} corresponding to ALP, collagen, and mineral fields. A basal production rate *r*_*b,i*_ applies to all live cells, while OBs contribute an additional osteogenic production rate *r*_*o,i*_ to the ALP, collagen, and mineral fields. No basal mineral production is included (*r*_*b,H*_ = 0).

Osteogenic supplementation is introduced at time *t*_osteo_ through the induction function

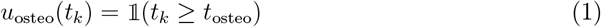

which activates osteogenic production terms.

Dex and MK-4 modulate osteogenic production through dimensionless response functions *m*_*i*_(*c*_*D*_, *c*_*K*_) acting on the osteogenic production term *r*_*o,i*_, which is co-induced with the osteogenic medium.

#### 2.4.2 Compound modulation

For each marker *i* ∈ {*A, C, H*} (ALP, collagen, HAp), Dex and MK-4 effects were modeled using biphasic log-dose response functions combined additively.

Log-dose offsets were defined as

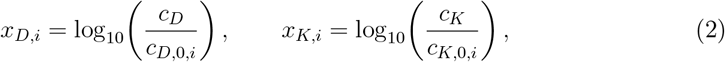

where *c*_*D*_ and *c*_*K*_ denote Dex and MK-4 concentrations, and *c*_*D*,0,*i*_, *c*_*K*,0,*i*_ are marker-specific reference concentrations associated with maximal response. For zero concentration, effectiveness was defined as *g*_*D,i*_ = 0 and *g*_*K,i*_ = 0.

Single-compound effectiveness functions were defined as clipped log-parabolic responses,

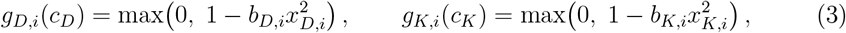

such that *g*_*D,i*_, *g*_*K,i*_ ∈ [0, 1] represent relative compound effectiveness in log-dose space. Parameters *b*_*D,i*_, *b*_*K,i*_ > 0 control the width of the biphasic response around the optimal concentration.

Combined modulation was assumed additive relative to the osteogenic baseline,

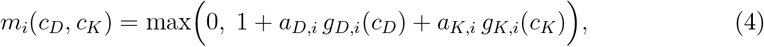

where *a*_*D,i*_, *a*_*K,i*_ > 0 specify the maximal contribution of Dex and MK-4 to osteogenic production for marker *i*. No explicit Dex–MK-4 interaction term was included.

#### 2.4.3 ALP dynamics

ALP activity evolves as

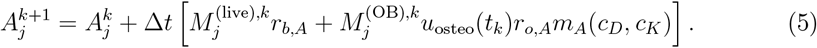

where *r*_*b,A*_ denotes the basal ALP production rate in live cells, and *r*_*o,A*_ denotes the additional osteogenic ALP production rate in OBs following osteogenic induction. The function *m*_*A*_(*c*_*D*_, *c*_*K*_) represents the dimensionless modulation factor describing the combined influence of Dex and MK-4 on ALP production.

The ALP field was initialized with baseline 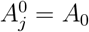, representing background media activity.

#### 2.4.4 Collagen deposition

The osteogenic component of collagen production was modeled as AA-dependent using a Michaelis–Menten-type factor.

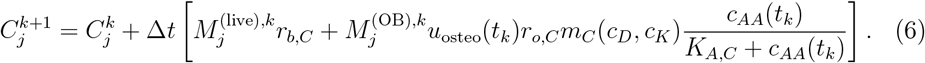

The initial condition is 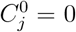. Here *r*_*b,C*_ denotes the basal collagen production rate in live cells, and *r*_*o,C*_ denotes the additional osteogenic collagen production rate in OBs after induction.

*c*_*AA*_(*t*_*k*_) denotes the AA concentration resulting from repeated medium supplementation and exponential decay, while *K*_*A,C*_ defines the half-saturation constant for AA-dependent collagen production.

The modulation factor *m*_*C*_(*c*_*D*_, *c*_*K*_) describes the combined effects of Dex and MK-4 on osteogenic collagen production.

#### 2.4.5 Mineralization dynamics

Mineralization occurs in OBs and depends on both extracellular matrix maturation and elapsed time since commitment. Commitment time is stored as a site-associated variable *t*_*j*,commit_ for lattice sites occupied by OBs. The effective post-commitment time is therefore defined as

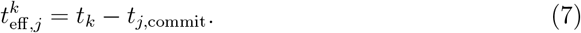

where *t*_*j*,commit_ denotes the time at which the OB currently occupying site *j* committed. Delayed matrix maturation is represented by a mature collagen field 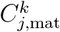, which relaxes toward the deposited collagen level with timescale *τ*_col_:

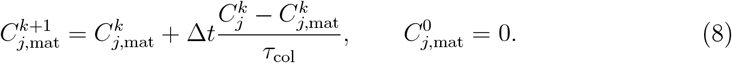

Matrix-supported mineralization capacity follows a saturating dependence on matured collagen,

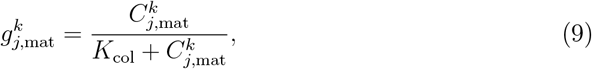

while commitment-dependent mineral competence follows Hill-type kinetics as a function of elapsed post-commitment time,

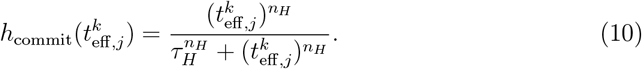

Mineralization evolves as

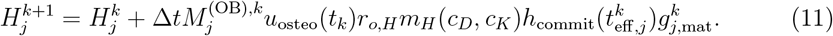

Here 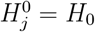 denotes detectable background mineral signal. Parameter *τ*_col_ controls collagen maturation, *K*_col_ defines the half-saturation level for matrix-supported mineralization, and (*τ*_*H*_, *n*_*H*_) determine the timing and steepness of post-commitment mineral competence.

### 2.5 Literature-derived parameterization

Model structure and parameter values were constrained from prior mechanistic studies rather than inferred from the endpoint dataset presented here. Parameters were assembled from literature describing osteoblast proliferation, differentiation timing, matrix production kinetics, and Dex/MK-4 dose responses. Marker production rates and compound modulation terms were estimated by fitting reduced submodels to published ALP, collagen, mineralization, and dose–response data. Details of the fitting procedures and resulting parameter estimates are provided in S1 Text, Section 2.

Parameters used in simulations are summarized in Table 1. Probabilities are defined per timestep (Δ*t* = 1 h).

**Table 1:** Model parameters used in OsteoMin simulations.

| Parameter | Symbol | Value | Description | Source |
| --- | --- | --- | --- | --- |
| Lattice spacing | $\Delta x$ | 30 $\mu\text{m}$ | Spatial resolution | [29] |
| Time step | $\Delta t$ | 1 h | Simulation timestep | chosen |
| Initial cell number | $N_0$ | 72 | Pre-OB count (S1 Text, Section 2.5) | experiment |
| Migration probability | $p_m$ | 0.27 $\text{h}^{-1}$ | Migration attempt probability | [29] |
| Division period | $\tau_i$ | $U(38, 52)$ h | Cell cycle duration | [30] |
| Commitment rate | $k_{\text{commit}}$ | 0.0125 $\text{h}^{-1}$ | OB differentiation rate | [31] |
| Maturation delay | $t_d$ | 72 h | Differentiation delay | [2, 3, 31] |
| Apoptosis probability | $p_a$ | $1 \times 10^{-3} \text{h}^{-1}$ | pre-OB death probability | [32] |
| AA half-saturation constant | $K_{A,C}$ | 5 $\mu\text{g mL}^{-1}$ | AA-dependent collagen gating | [34] |
| Ascorbate half-life | $t_{1/2}^{AA}$ | 8 h | AA degradation in medium | [34] |
| Collagen half-saturation | $K_{\text{col}}$ | 1.0 | Matrix for half-maximal deposition | [12] |
| Collagen maturation time | $\tau_{\text{col}}$ | 48 h | ECM maturation timescale | [38] |
| Mineral activation time | $\tau_H$ | 384 h | Mineralization onset timescale | [39] |
| Mineral Hill exponent | $n_H$ | 2 | Activation steepness | [39] |
| Basal ALP production | $r_{b,A}$ | $9.49 \times 10^{-2} \text{AU h}^{-1}$ | Baseline ALP synthesis | [40] |
| Osteogenic ALP production | $r_{o,A}$ | $8.10 \times 10^{-1} \text{AU h}^{-1}$ | OB-specific ALP synthesis | [40] |
| Basal collagen production | $r_{b,C}$ | $3.36 \times 10^{-3} \text{AU h}^{-1}$ | Baseline collagen synthesis | [38] |
| Osteogenic collagen production | $r_{o,C}$ | $1.30 \times 10^{-2} \text{AU h}^{-1}$ | OB-specific collagen synthesis | [38] |
| Osteogenic mineral production | $r_{o,H}$ | $1.02 \times 10^{-1} \text{AU h}^{-1}$ | OB-specific mineralization | [39] |

A representative baseline simulation was performed over 28 days on a periodic 33 × 33 lattice with randomly seeded cells and no Dex or MK-4 perturbation. Given the assumed MC3T3-E1 cell footprint (Δ*x* ≈ 30 µm), this domain corresponds to an approximate culture area of 1 × 1 mm, comparable to the spatial scale of a typical microscopy field of view. Cells undergo stochastic migration, pre-OB proliferation, apoptosis, and delayed osteogenic commitment following induction. ALP, collagen matrix, and mineral fields are produced locally and accumulate over time. Representative model outputs at assay-relevant times are shown for cell-state distribution (Fig. 2a), ALP production (Fig. 2b), and collagen–mineral co-localization (Fig. 2c).

**Fig 2:**
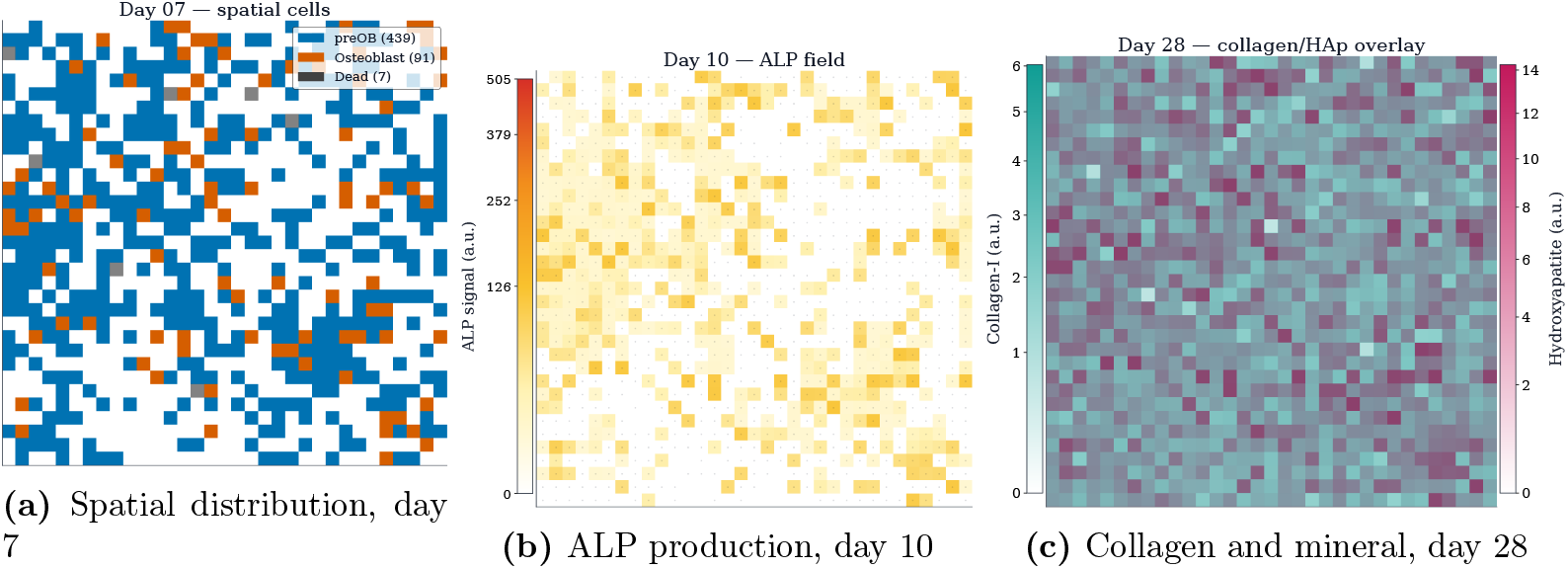
Spatiotemporal progression of osteogenic differentiation in OsteoMin. (A) Cell distribution at day 7. (B) ALP field at day 10. (C) Collagen and mineral fields at day 28.

### 2.6 Statistical Analysis

All experimental endpoint data are presented as individual points. Group differences were evaluated using the non-parametric Kruskal–Wallis test with two-sided Mann–Whitney U post hoc comparisons when significant. P-values were adjusted using the Benjamini– Hochberg procedure (false discovery rate control). Adjusted p-values are reported as q-values, with significance defined as *q* < 0.05 (*) and *q* < 0.01 (**).

Analyses were chosen according to the diagnostic objective of each model experiment. Concordance between simulated and experimental relative responses was evaluated using Kendall’s *τ* and directional agreement relative to the osteogenic condition. Separability of perturbation families was quantified using leave-one-out 1-nearest-neighbor classification of marker-specific trajectories (Euclidean distance), summarized by macro-F1. Parameter influence on assay-aligned endpoints was assessed using variance-based Sobol indices with Saltelli sampling (SALib).

### 2.7 Implementation and Performance

The model was implemented in Python. Simulations ran on an AMD Ryzen 7 7435HS CPU (3.10 GHz), NVIDIA RTX 4060 Laptop GPU, and 16 GB RAM. Simulations used a 33 × 33 lattice (1,089 sites) seeded with 72 cells and run for 672 biological hours (28 days). Each 28-day simulation required approximately 22 seconds on a single CPU core (seed = 12345).

## 3 Experimental Methods

MC3T3-E1 pre-OB cells were cultured under basal and systematically varied osteogenic conditions to assess differentiation and mineralization responses.

### 3.1 Cell Culture and Osteogenic Induction

MC3T3-E1 Subclone 4 cells were cultured in *α*-MEM (no ascorbic acid) supplemented with 10% FBS and 1% Antibiotic–Antimycotic at 37 °C and 5% CO_2_. Cell concentration was determined by hemocytometer counting, yielding ~ 4.1 × 10^5^ cells/mL. For experiments, 12 mm glass coverslips (area ≈ 113 mm^2^) in 24-well plates were seeded at 8.0 × 10^3^ cells/well (density ≈ 72 cells/mm^2^) in 0.4 mL medium. Osteogenic supplements (AA, *β*-GP, Dex, MK-4) were added on Day 3, with medium changes every 48–72 h. The medium compositions are listed in Table 2. Detailed reagent information, including suppliers and catalog numbers, is provided in S1 Text, Section 6.

**Table 2.** Experimental media compositions. All conditions contain ethanol carrier (0.10 % v/v).

| ID | Condition | AA (mM) | β-GP (mM) | Dex (nM) | MK-4 (μM) |
| --- | --- | --- | --- | --- | --- |
| C1 | Basal | 0 | 0 | 0 | 0 |
| C2 | AA + β-GP (Osteo) | 0.284 | 10 | 0 | 0 |
| C3 | Osteo + $D_L K_L$ | 0.284 | 10 | 10 | 0.1 |
| C4 | Osteo + $D_L K_H$ | 0.284 | 10 | 10 | 10 |
| C5 | Osteo + $D_H K_L$ | 0.284 | 10 | 100 | 0.1 |
| C6 | Osteo + $D_H K_H$ | 0.284 | 10 | 100 | 10 |
| C7 | Osteo + $D_H K_0$ | 0.284 | 10 | 100 | 0 |
| C8 | Osteo + $D_0 K_H$ | 0.284 | 10 | 0 | 10 |

### 3.2 Endpoint Assays

ALP activity, collagen type I immunofluorescence, and Alizarin Red staining were performed on days 10, 14, and 28, respectively (n=3 biological replicates). These time points reflect early differentiation, matrix maturation, and late mineralization.

In addition to endpoint measurements, brightfield images were acquired using an Olympus IX81 inverted microscope with an sMAX04BM sCMOS camera and 4× and 10× objectives to monitor cell growth and mineralization throughout culture. Representative images are provided in S1 Text, Section 5.

#### 3.2.1 Alkaline Phosphatase Activity

Intracellular ALP activity was assessed on day 10. Culture medium was aspirated, and cells were rinsed once with ice-cold Ca^2+^/Mg^2+^-free DPBS to remove residual serum components. Cells were lysed using an ice-cold, phosphate-free buffer composed of 50 mM Tris, 1 mM MgCl_2_, and 0.2% Triton X-100. Lysates were clarified by centrifugation at 10,000 × *g* for 10 min at 4 °C, aliquoted, and snap-frozen up to analysis time.

For enzymatic quantification, 25 µL of lysate was incubated with 75 µL of SIGMAFAST^™^ p-nitrophenyl phosphate (pNPP) substrate solution for 30 min at room temperature. The reaction was terminated by the addition of 25 µL of 3 M NaOH, and absorbance was measured at 405 nm. Absorbance values were converted to ALP activity (mU· mL−1) assuming linear proportionality to enzymatic rate, using calf intestinal phosphatase as reference.

#### 3.2.2 Collagen type I Immunofluorescence

Extracellular matrix deposition and cellular organization were assessed by immunofluorescence staining of collagen type I, nuclei, and F-actin. Cells were rinsed with pre-warmed Ca^2+^/Mg^2+^-free DPBS (pH 7.4) and fixed with freshly prepared 4% paraformaldehyde in DPBS for 15 min at room temperature, followed by three DPBS washes.

Cells were permeabilized with 0.1% Triton X-100 in DPBS for 5 min and blocked with 1% bovine serum albumin (BSA) in DPBS for 60 min at room temperature. Samples were incubated overnight at 4 °C with mouse anti-collagen type I primary antibody (1:300 in 1% BSA/DPBS).

After overnight incubation, samples were washed and incubated for 60 min at room temperature in the dark with goat anti-mouse Alexa Fluor 488 secondary antibody (1:500) and phalloidin–Alexa Fluor 568 (1:500). Nuclei were stained with DAPI (300 nM) for 15 min. Coverslips were mounted using Mowiol mounting medium and imaged the same day.

Images were acquired using a Leica SP8 confocal microscope equipped with an HC PL APO CS2 20×/0.75 IMM objective. Images of DAPI, actin, and collagen channels were acquired sequentially from 5 ROIs for all experimental conditions to allow quantitative comparison.

#### 3.2.3 Alizarin Red S Staining

Matrix mineralization was assessed on day 28 using ARS staining. Cultures in 24-well plates were rinsed with PBS and fixed with 4% paraformaldehyde in PBS for 15 min at room temperature. The ARS working solution was prepared at 1% (w/v), adjusted to pH 4.15 using 0.1 M NaOH, and filtered through a 0.22 µm membrane to ensure selective binding to calcium-containing deposits. Fixed samples were incubated with the ARS solution for 15 min at room temperature under gentle agitation in the dark, followed by five washes with deionized water until complete removal of unbound dye, and then imaged.

Brightfield images were acquired using an Olympus IX-PHL inverted microscope equipped with a UPLFLN 4×/0.13 PH1 objective and an Olympus DP71 CCD camera. Nine fields of view were imaged per condition using identical exposure settings.

Mineralization was quantified as the fraction of ARS-positive pixels relative to total image area [41]. Images were processed using automated thresholding based on the complementary color ratio *R/*(*G*+*B*), where *R, G*, and *B* denote red, green, and blue channel intensities. Pixels with *R/*(*G* + *B*) ≥ 0.55 were classified as ARS-positive. Threshold sensitivity analysis is provided in S1 Text, Section 4.

## 4 Results

### 4.1 Experimental assay readouts

Osteogenic progression was evaluated at days 10, 14, and 28, corresponding to early enzymatic activity, extracellular matrix deposition, and late-stage mineralization.

#### 4.1.1 ALP activity

ALP activity increased under osteogenic conditions relative to basal culture (Fig. 3A). Dex and MK-4 supplementation produced variable responses without a consistent monotonic dose dependence. Differences between supplemented conditions were modest relative to replicate variability, and no statistically significant differences were observed.

**Fig 3:**
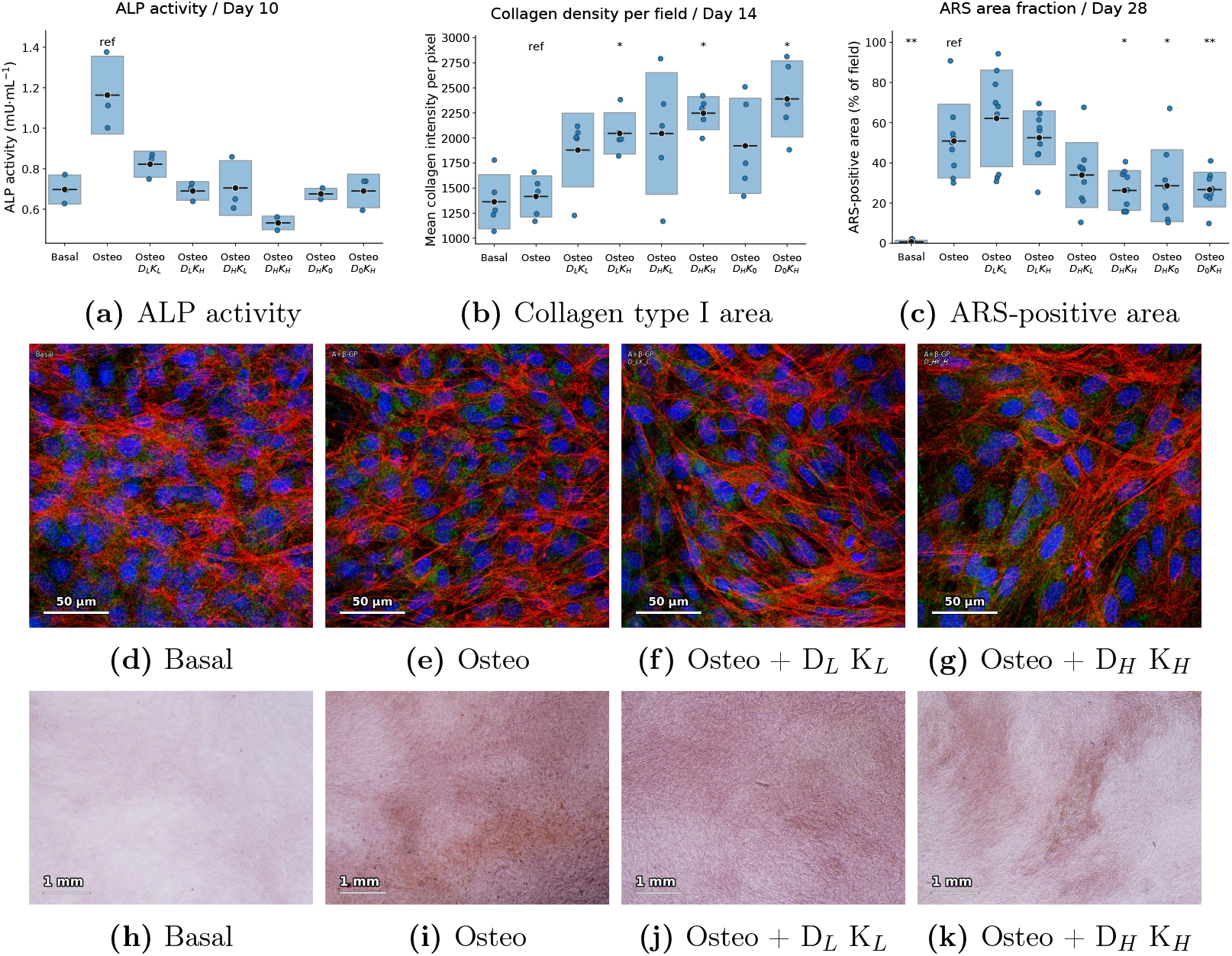
Experimental osteogenic differentiation readouts across culture conditions. Quantification of ALP activity (day 10), collagen type I area fraction (day 14), and ARS-positive area fraction (day 28) are shown in (A-C). Representative confocal images of nuclei (DAPI), F-actin, and collagen type I are shown in (D-G). Representative ARS images of mineralized matrix are shown in (H-K).

#### 4.1.2 Collagen deposition

Collagen type I area increased under osteogenic conditions compared to basal culture (Fig. 3B). MK-4-containing conditions showed a modest increase relative to osteogenic medium alone, with substantial overlap between conditions.

Confocal imaging showed sparse collagen signal and less organized cytoskeletal structure in basal cultures (Fig. 3D). Osteogenic conditions exhibited increased cell spreading and denser collagen networks (Fig. 3E). Dex and MK-4 supplementation was associated with increased collagen signal intensity (Fig. 3F-G) and organized actin filaments.

#### 4.1.3 Mineralized area

Mineralized area fraction increased under osteogenic conditions relative to basal culture (Fig. 3C). Differences between Dex/MK-4 conditions were moderate, with low Dex and MK-4 concentrations associated with increased mineralization.

ARS staining showed heterogeneous mineralization with localized nodular regions (Fig. 3H-K). Spatial heterogeneity was further characterized using local variance analysis (S1 Text, Section 5).

### 4.2 OsteoMin trajectories and module ablation

We first examined baseline temporal behavior and then quantified the contribution of individual mechanisms by selective module ablation. We use ablation in the computational sense: the model is rerun with a specific component disabled (e.g., *k*_commit_ = 0, no apoptosis, no delay, or removal of the 2D spatial representation) and compared to the full model.

The reference spatial model generated assay-aligned trajectories for ALP, collagen, and mineralization (Fig. 4A). ALP increased toward day 10, collagen increased toward day 14, and mineralization showed delayed onset with stronger growth toward day 28.

**Fig 4:**
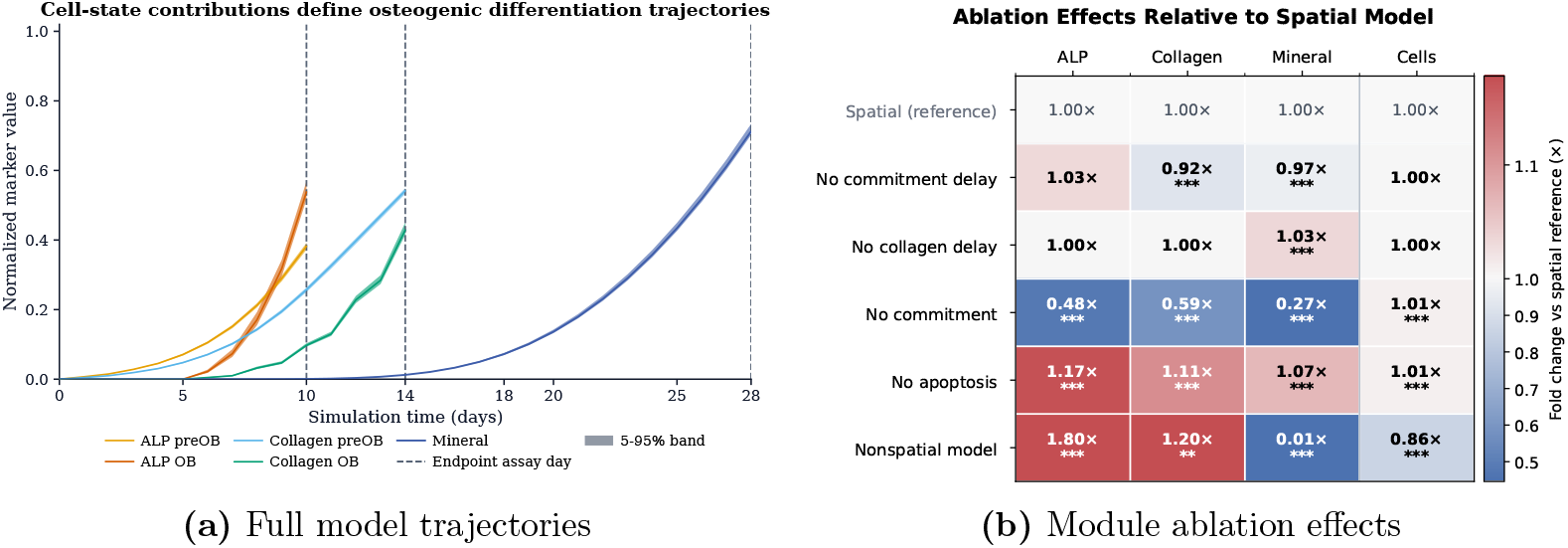
Baseline OsteoMin trajectories and module ablation analysis. (A) Simulated time courses of ALP activity, collagen deposition, and mineralization in the reference model, with vertical dashed lines indicating assay-aligned readout times (days 10, 14, and 28). (B) Ablation effects at assay endpoints relative to the spatial reference model, shown as median fold change relative to the spatial reference model across independent simulations.

**Fig 5:**
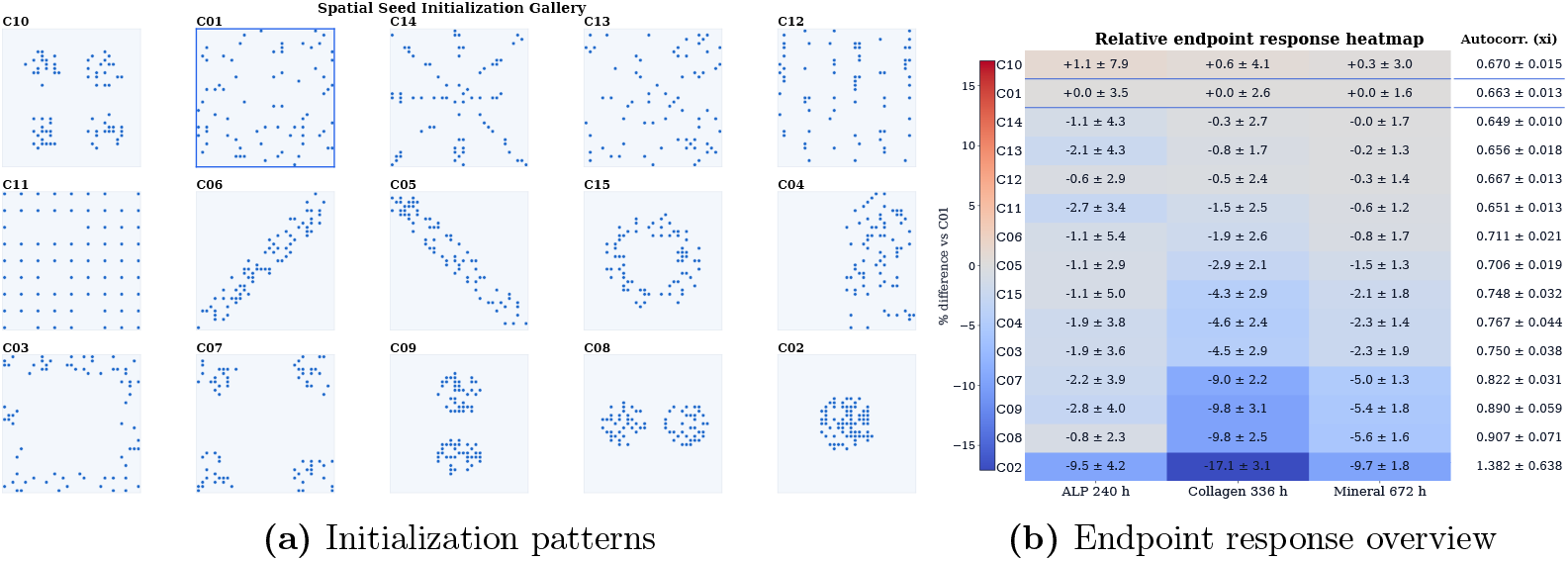
Initialization effects in the OsteoMin model. (A) Representative initial seeding patterns on the simulation lattice. (B) Percent differences in simulated ALP (240 h), collagen (336 h), and mineral (672 h) relative to the random-seeding control (C01). The left colorbar applies to these three heatmap endpoints only. The right column reports raw mineral-field autocorrelation length, at 672 h.

To assess mechanism-level contributions, each ablation was compared with the spatial reference model under identical baseline conditions. Nonspatial model details are provided in S1 Text, Section 3.2.1. Effects are reported as median fold change relative to the spatial reference model at assay endpoints across independent simulations, with significance from two-sided Mann–Whitney U tests and Benjamini–Hochberg correction (Fig. 4B).

Module ablation showed distinct endpoint effects that clarify the functional role of individual mechanisms. Removing lineage commitment strongly reduced ALP (0.48×), collagen (0.59×), and mineral (0.27×), indicating that commitment is the primary source of osteoblast-dependent matrix production and mineral formation. Removing apoptosis increased all readouts (ALP 1.17×, collagen 1.11×, mineral 1.07×), indicating that turnover primarily limits cumulative matrix accumulation rather than regulating differentiation timing. Removing commitment delay produced minimal change in ALP (1.03×) and final cell number (1.00×), but modest reductions in collagen (0.92×) and mineral (0.97×), suggesting that differentiation timing influences matrix maturation windows more than early marker expression. Eliminating collagen-maturation delay left ALP and collagen unchanged (1.00×) while slightly increasing mineral (1.03×), consistent with mineral formation being constrained by matrix readiness rather than matrix abundance. Relative to the spatial reference, the nonspatial comparator increased early markers (ALP 1.80×, collagen 1.20×) but markedly reduced mineralization (0.007×) and modestly reduced final cell number (0.86×). This divergence indicates that spatial organization primarily affects late-stage mineral emergence: without spatial co-localization constraints, matrix accumulates rapidly but fails to achieve the local density and maturation coupling required for nucleation-like behavior. Together, these results indicate that commitment governs osteogenic capacity, apoptosis regulates cumulative production, and spatial exclusion with heterogeneous matrix maturation generates threshold-like mineralization dynamics that are not captured by population-averaged formulations.

### 4.3 Latent trajectories underlying similar endpoints

To examine how differences in initial cell organization affect model behavior, we simulated multiple initial seeding patterns on the same periodic (1 × 1 mm) lattice while keeping all biological parameters constant. All initialization patterns used the same total number of initial cells (*N*_0_ = 72), so differences reflect spatial organization rather than initial population size or global density. Across the initialization panel, seeding ranged from uniform random placement (C01) to structured motifs, including compact center clustering (C02), edge-biased and half-domain layouts (C03–C04), diagonal bands (C05– C06), corner clusters and multi-lobe patterns (C07–C10), lattice, stripe, and checkerboard geometries (C11–C13), radial spokes (C14), and a peripheral ring (C15).

For each configuration, ALP activity, collagen deposition, and mineralization were quantified at the corresponding assay endpoints and compared with a uniform random seeding control (C01). Mineral-field autocorrelation at 672 h was quantified using two-point correlation analysis; details of the spatial autocorrelation metrics are provided in S1 Text, Section 4.

Relative to the reference condition (C01), endpoint variation was modest for ALP and mineralization but larger for collagen: ALP at 240 h ranged from +1.1% to −9.5%, collagen at 336 h from +0.6% to −17.1%, and mineral at 672 h from +0.3% to −9.7%. In particular, the central-cluster initialization (C02) showed the strongest reduction in mid-stage collagen accumulation. C10 slightly exceeded C01 because its quadrilobular seeding improved per-cell yield despite similar OB counts. Autocorrelation is shown as raw ξ: values span approximately 0.649 ± 0.010 to 1.382 ± 0.638, with C02 exhibiting the largest spatial correlation length. Overall, distinct spatial organizations can yield similar endpoint outcomes, indicating that endpoint assays underconstrain spatial aspects of differentiation and may obscure an important source of variability across experiments.

### 4.4 Effects of Dex and MK-4 on osteogenic markers

Model responses to dexamethasone (Dex) and menaquinone-4 (MK-4) were evaluated across literature-reported concentration ranges (0–100 nM for Dex; 0–10 µM for MK-4) using single-factor parameter sweeps. Discrete dose grids are described in S1 Text, Section 3.2.

Endpoint values were extracted at assay-aligned time points (ALP: 240 h; collagen: 336 h; mineral: 672 h) and expressed as log_2_ fold change relative to the osteogenic reference condition C2 (Dex = 0, MK-4 = 0).

To evaluate combined effects, Dex and MK-4 concentrations were varied jointly over a 10 × 10 grid spanning the same concentration ranges, yielding 100 conditions. For each condition, endpoint responses were computed using the same normalization procedure.

Dex single-factor responses exhibited biphasic behavior, with increases up to intermediate concentrations followed by mild declines at higher doses. ALP peaked at approximately 30 nM, collagen at approximately 20 nM, and mineral at approximately 60 nM, with only marginal changes observed between 60 and 100 nM. MK-4 peak windows were ~0.6–1 µM for ALP and mineralization (maximum at 0.6 µM), and 0.2–0.6 µM for collagen (maximum at 0.3 µM) (Fig. 6A).

**Fig 6:**
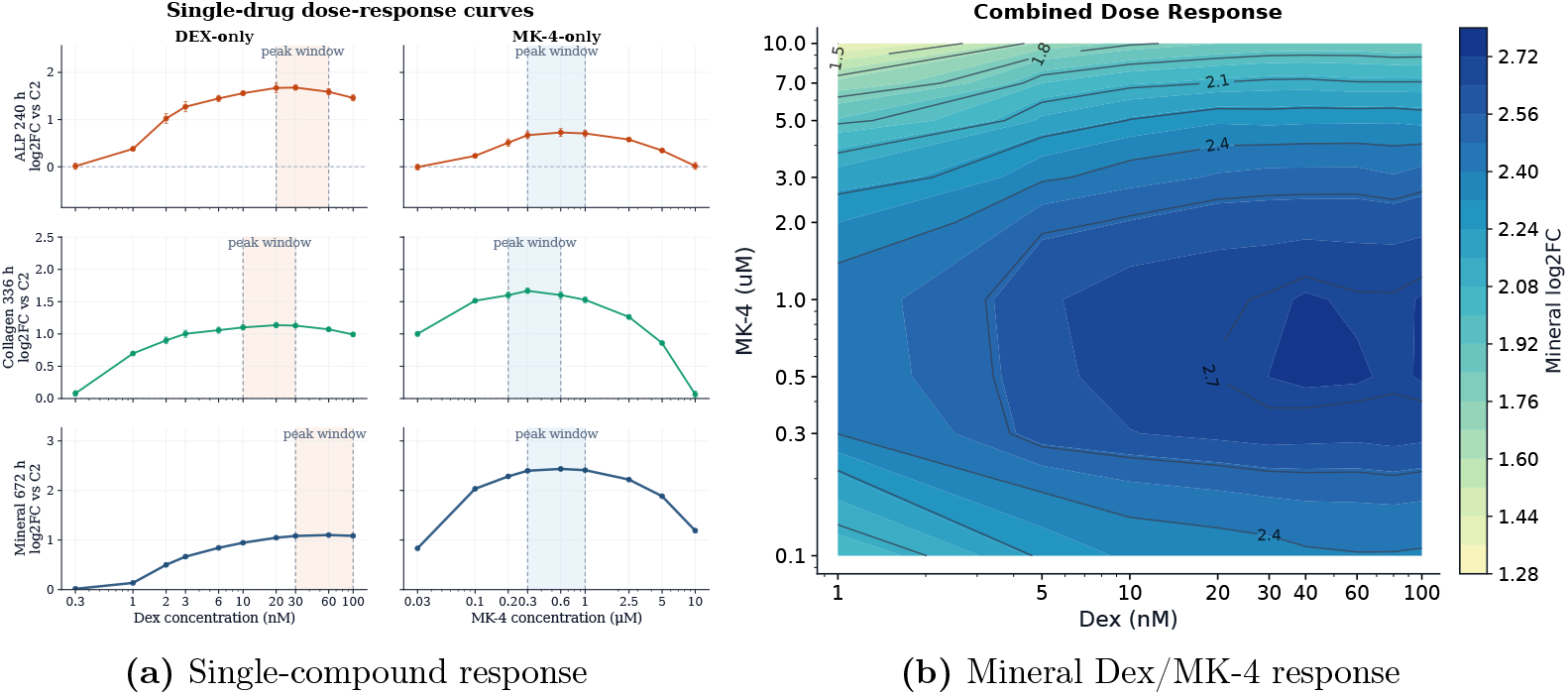
Dose-response analysis of model predictions. (A) Single-compound dose-response curves for Dex and MK-4 showing ALP (240 h), collagen (336 h), and mineral (672 h) as log_2_ fold change versus C2. (B) Combined Dex–MK-4 response surface summarized as log_2_ fold change versus C2 across the two-dimensional concentration grid.

**Fig 7:**
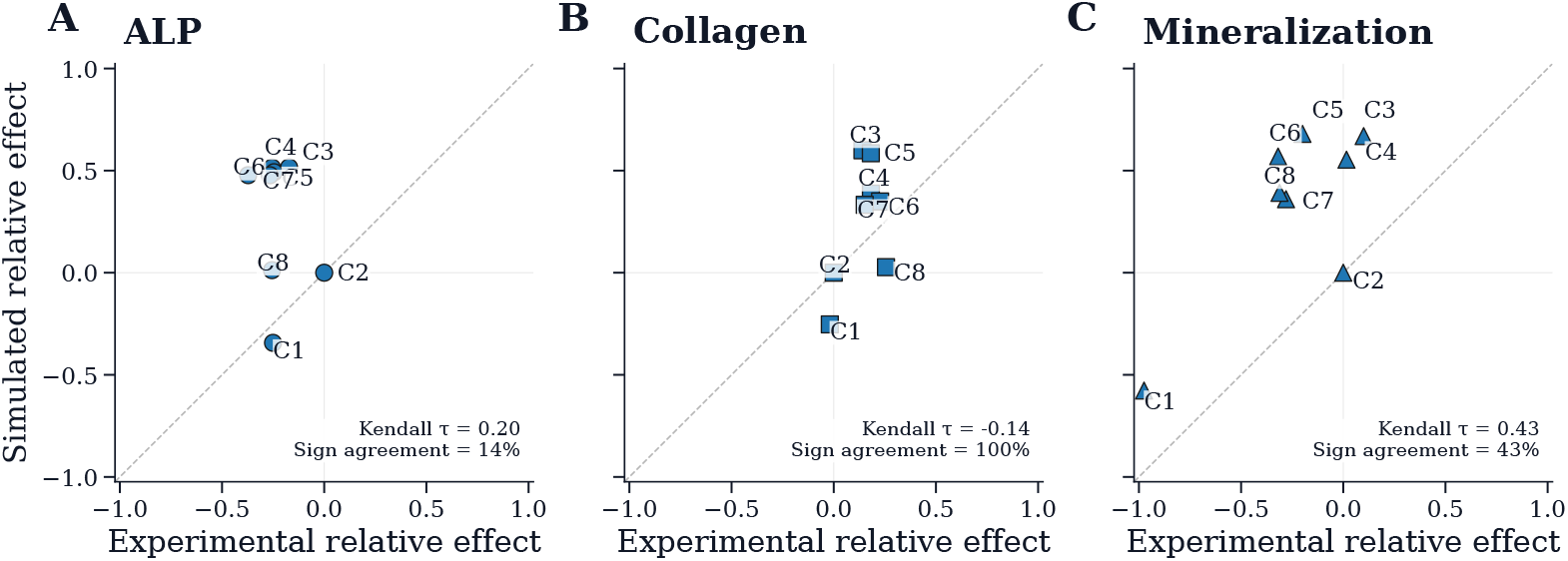
Model–experiment comparison for normalized osteogenic outcomes. Simulated and experimental relative responses, normalized to C2, are shown for ALP (A), collagen (B), and mineralization (C). Concordance across the seven non-reference conditions is summarized by Kendall’s *τ*, and sign agreement.

Combined responses were strongest within a bounded Dex–MK-4 region rather than at uniformly maximal single-factor doses (Fig. 6B). In the two-dimensional mineral response surface (log_2_ fold change vs baseline osteogenic C2 condition), high responses form a broad plateau at intermediate-to-high Dex (~20–100 nM) with low-to-moderate MK-4 (~0.3–2 µM), while responses decline at high MK-4 (≳7–10 µM) across Dex levels. The local maximum is observed near Dex ~40–60 nM and MK-4 ~0.5–1 µM. Overall, the landscape indicates that multiple Dex–MK-4 pairs can achieve similarly high mineral outcomes within non-toxic concentration ranges.

### 4.5 Comparison of model and experimental endpoint responses

To evaluate agreement between independently obtained experimental measurements and literature-parameterized simulations, ALP, collagen, and mineralization endpoints were compared across the eight media conditions in Table 2. Model parameters were derived from prior studies and were not fitted to the experimental dataset presented here. The comparison therefore serves as an external consistency check of whether literature-supported differentiation dynamics produce qualitatively consistent endpoint trends under commonly used perturbations.

Responses were compared using a reference-centered effect metric relative to the osteogenic reference condition (C2):

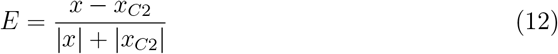

where *x* denotes the endpoint value under a given condition and *x*_*C*2_ is the corresponding value under the reference osteogenic condition.

Concordance between simulated and experimental relative responses was evaluated using Kendall rank correlation (*τ*) and directional agreement (sign concordance) across non-reference conditions (*n* = 7). Kendall *τ* quantifies consistency in condition ordering, while sign agreement measures concordance in response direction relative to the reference condition.

ALP showed limited directional agreement (14%) with weak rank concordance (*τ* = 0.20). Collagen showed full directional agreement (100%) but negative rank concordance (*τ* = −0.14), indicating consistent response direction but poor ordering agreement across conditions. Mineralization (ARS) showed moderate directional agreement (43%) with moderate rank concordance (*τ* = 0.43). Disagreement was concentrated in high-Dex conditions (C5–C7; Dex = 100 nM), especially for ALP and ARS, where simulated relative effects remained above C2 while experimental effects were reduced.

Overall, the comparison showed partial consistency but also substantial marker-specific discrepancies, especially under high-Dex conditions. This pattern supports the interpretation that endpoint agreement alone is insufficient to uniquely constrain the underlying mechanisms.

### 4.6 Endpoint observations weakly identify latent perturbation mechanisms

We evaluated whether staged endpoint assays contain sufficient information to distinguish mechanistic perturbation families arising from subclonal variation, spatial initialization, differentiation dynamics, and media-dependent modulation. These sources of variability are commonly associated with reproducibility differences in osteogenic experiments. Parameter ranges and sampling schemes for all perturbation families are provided in S1 Text, Section 3.3.

We considered four perturbation families: spatial initialization, cell number, differentiation dynamics, and ascorbic-acid–mediated media effects, each comprising 50 parameterized conditions with 10 stochastic replicates. Replicate means were used to construct marker-specific endpoint and trajectory feature vectors. Values were standardized across conditions (for trajectories, per day), and Euclidean distances were used for leave-one-out nearest-neighbor classification of perturbation families. Performance was quantified using macro-F1, defined as the unweighted mean of per-family F1 scores.

Let *z*(*t*; *θ, x*_0_) denote the latent differentiation trajectory determined by model parameters *θ* and spatial initialization *x*_0_. Experimentally accessible measurements correspond to discrete observations of marker dynamics at assay-aligned timepoints (days 10, 14, and 28). For each marker *m* ∈ {ALP, Collagen, Mineral} and condition *i*, we define (i) an endpoint feature and (ii) a truncated trajectory feature:

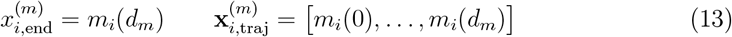

with (*d*_ALP_, *d*_Collagen_, *d*_Mineral_) = (10, 14, 28).

We emphasize that identifiability is assessed within the model class and observation scheme considered here. The results therefore quantify practical separability of mechanistic hypotheses under endpoint sampling, rather than intrinsic biological identifiability.

Macro-F1 ranges from 0 to 1, with a random baseline of 0.25 for this four-class problem. In the split-cell matrices (Fig. 8), trajectory-based classifications show consistently stronger diagonal dominance than endpoint-based classifications, indicating improved recovery of true perturbation families. Endpoint scores remain modest (*F* 1_ALP@10_ = 0.49, *F* 1_Collagen@14_ = 0.40, *F* 1_Mineral@28_ = 0.39), whereas trajectory-based scores are substantially higher (*F* 1_ALP(0–10)_ = 0.70, *F* 1_Collagen(0–14)_ = 0.86, *F* 1_Mineral(0–28)_ = 0.79).

**Fig 8:**
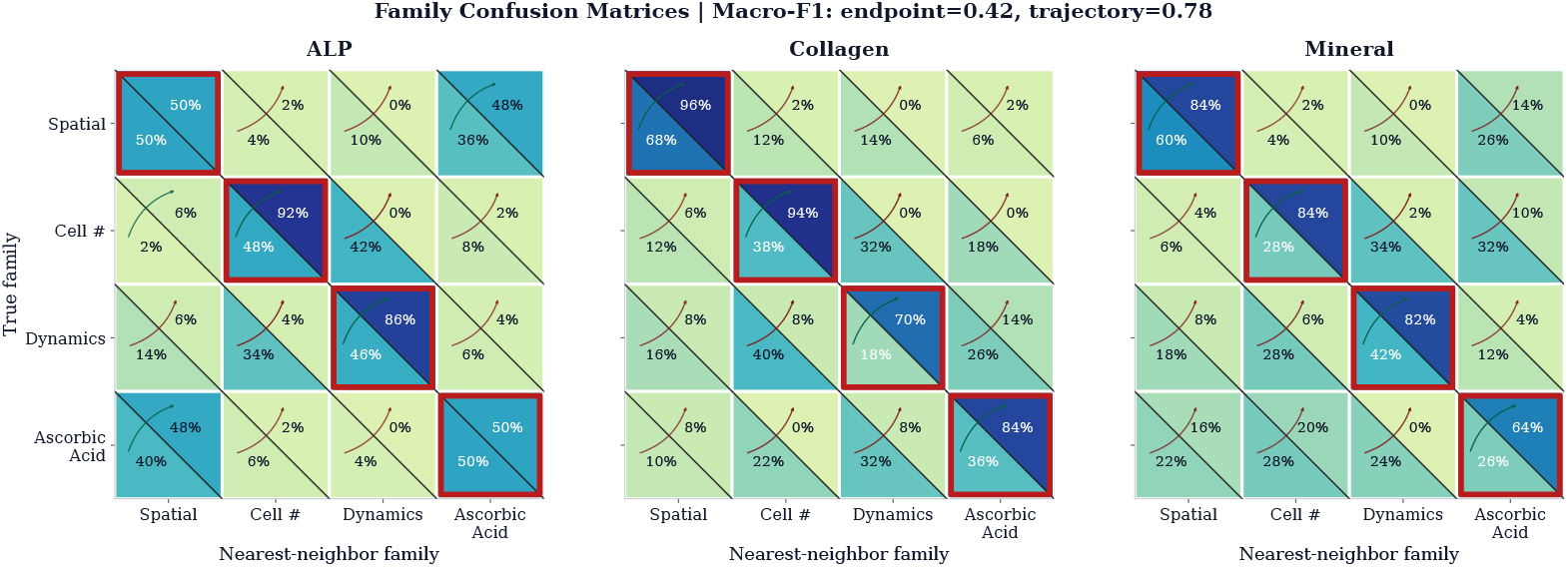
Classification of perturbation families from marker-specific readouts using split-cell confusion matrices. Lower-left triangles show endpoint-based classification (ALP day 10, collagen day 14, mineral day 28), and upper-right triangles show trajectory-based classification over the corresponding intervals. Red-outlined squares denote true positives (correct family classification). Across markers, endpoint readouts exhibit low separability (macro-F1 ≈ 0.42, 95% CI 0.39–0.46), whereas trajectories improve separability (macro-F1 ≈ 0.78, 95% CI 0.75–0.81). This demonstrates a many-to-one mapping from latent perturbations to endpoint observables, indicating structural non-identifiability under endpoint sampling, which is partially resolved by temporal trajectories.

This pattern reflects a collapse of distinct mechanistic perturbations onto similar endpoint observables, consistent with a many-to-one mapping from latent trajectories to measured outputs. Incorporating temporal information partially lifts this degeneracy, enabling improved discrimination between perturbation families.

### 4.7 Global Sensitivity Analysis identifies dominant regulatory parameters

Global sensitivity analysis was performed using variance-based Sobol indices implemented in SALib [42] to quantify parameter influence on simulated osteogenic endpoints. A total of *D* = 20 model parameter ranges were varied jointly across biologically plausible ranges (S1 Text, Section 3) using Saltelli’s extension of Sobol quasi-random sampling with base sample size *N* = 256, without computing second-order indices, resulting in *N* (*D* + 2) = 5,632 model evaluations. The sampled parameter sets were drawn from uniform distributions over the specified bounds. Endpoints were extracted at assay-aligned timepoints (ALP 240 h, collagen 336 h, mineral 672 h).

Sensitivity results are summarized in Fig. 9. Panels A–C show first-order and total-order indices for ALP, collagen, and mineral endpoints. Error bars indicate 95% confidence intervals.

**Fig 9:**
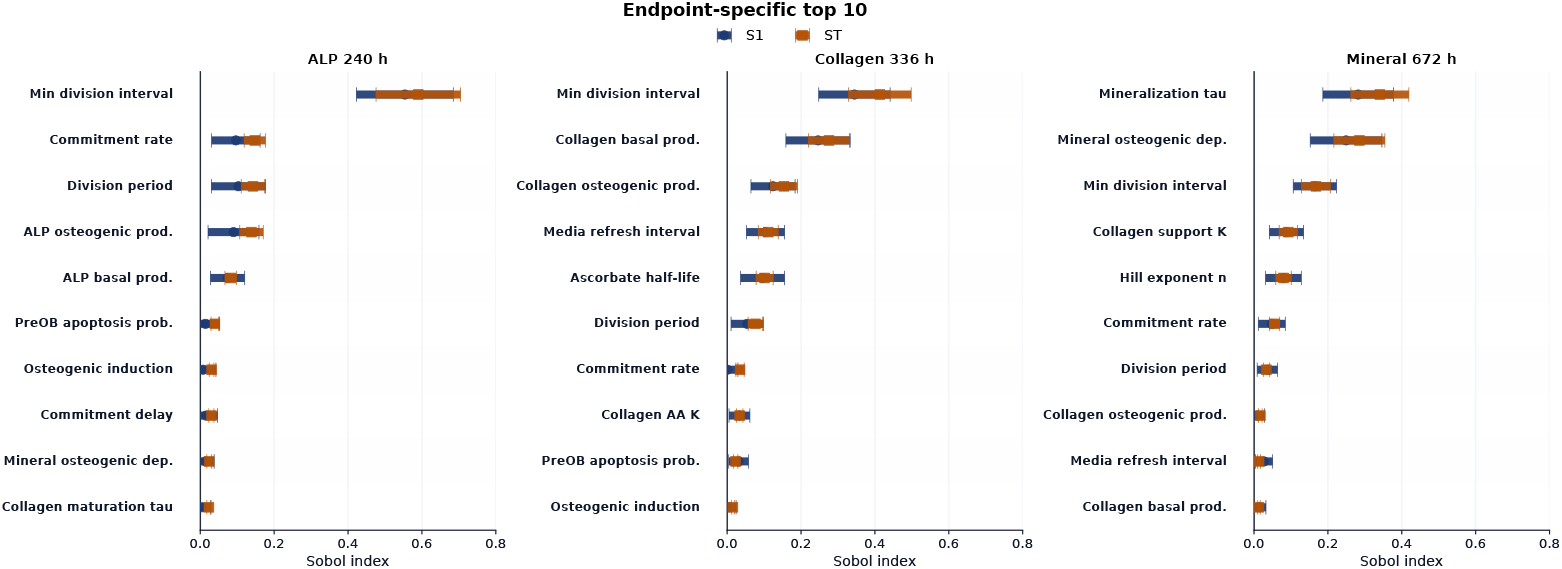
Sobol sensitivity analysis of model parameters. First-order (*S*_1_) and total-order (*S*_*T*_) Sobol indices are shown for (A) ALP activity at day 10, (B) collagen deposition at day 14, and (C) mineralization at day 28, indicating the individual and overall contributions of each parameter to output variance.

Sensitivity analysis showed distinct parameter influence across the differentiation program. Early ALP levels were primarily influenced by proliferation timing, with the minimum division interval contributing the largest variance (*S*_*T*_ ≈ 0.62), consistent with ALP reflecting both lineage progression and population expansion [3, 39]. Collagen deposition depended on both division dynamics and basal and osteogenic collagen production, indicating combined effects of cell number and per-cell matrix secretion [7, 38]. Late mineralization was dominated by nonlinear mineral-specific parameters, particularly the Hill exponent governing mineral competence (*S*_*T*_ ≈ 0.41), consistent with delayed matrix maturation preceding mineralization [38, 43]. The larger difference between first- and total-order indices for mineralization indicates interaction effects, suggesting that late-stage outputs depend on coupled upstream processes. Overall, different assay endpoints probe distinct components of the differentiation program.

### 4.8 Generalizability of the framework

Although OsteoMin was developed for osteogenic differentiation, the framework applies more broadly to in vitro systems in which cells undergo discrete state transitions, interact in space, and are primarily characterized through aggregate population-level measurements. In such settings, observable outputs reflect heterogeneous population behavior, while intermediate progression remains only partially resolved. Consequently, multiple latent dynamics may be consistent with similar endpoint readouts.

Rather than uniquely identifying mechanisms from endpoint data alone, the framework defines a constrained space of admissible latent dynamics. Given initial conditions and candidate transition rates, the model can be propagated forward to generate experimentally testable predictions across perturbations, doses, and sampling schemes. Comparison with independent measurements progressively excludes incompatible parameter regimes and supports hypothesis discrimination and experimental design in incompletely observed systems.

## 5 Discussion

OsteoMin treats ALP activity, collagen deposition, and mineralization as partial observations of an underlying stochastic cell-state process. Within this framework, endpoint assays correspond to an incomplete projection of differentiation dynamics, such that distinct mechanistic pathways—arising from different kinetic parameters or spatial initial conditions—can produce similar observable outcomes. This reflects a general limitation of inferring system dynamics from sparsely sampled, population-averaged measurements [44].

This limitation is supported quantitatively by the model results. Using endpoint measurements alone, alternative mechanistic hypotheses were only weakly separable (macro-F1 ≈ 0.42), whereas incorporating temporal trajectories substantially improved discrimination (macro-F1 ≈ 0.78). This indicates that trajectory information, rather than terminal marker values alone, carries much of the mechanistic signal.

Spatial initialization further influences assay outcomes. The spatial formulation is motivated by the dependence of matrix accumulation and mineral nucleation on local cell density and neighborhood history, which are not directly represented in mean-field models without additional assumptions. Despite substantial variation in spatial organi-zation, endpoint differences remained limited (<20% for collagen and ~10% for mineral), indicating that spatial heterogeneity is only weakly reflected in standard assay readouts [3, 43]. In the model, mineralization emerges from locally coupled matrix maturation and deposition, producing heterogeneous and clustered patterns consistent with experimental observations [45]. However, these differences are largely obscured when aggregated to endpoint measurements, such that distinct spatial organizations yield similar assay-level outcomes.

Compound response analysis shows that the model reproduces biphasic dexamethasone responses, with intermediate concentrations associated with increased osteogenic markers [35, 46, 47]. At the same time, multiple Dex/MK-4 combinations produce comparable endpoint responses, indicating that different regulatory regimes can map to similar observable states. This highlights that agreement with endpoint data alone does not uniquely constrain the underlying mechanism, even when model assumptions are consistent with known biology. Discrepancies at high Dex concentrations, where experimental inhibition is not reproduced, may reflect additional processes such as mitochondrial dysfunction and apoptosis [48], as well as the dependence of ALP measurements on sampling time [49]. Including these effects would introduce additional poorly constrained parameters, increasing model flexibility without resolving the underlying identifiability limitations.

Sensitivity analysis indicates that parameter influence shifts across the differentiation program. Early ALP variability is primarily associated with proliferation timing (*S*_*T*_ ≈ 0.62), whereas late mineralization depends on nonlinear parameters governing matrix competence (*S*_*T*_ ≈ 0.41). The larger difference between first- and total-order indices for mineralization suggests interaction effects, implying that late-stage outcomes depend on coupled upstream processes. As a result, endpoint measurements at a single time point provide limited ability to disentangle parameter contributions, further constraining practical identifiability.

Several limitations should be noted. OsteoMin was designed as a coarse-grained model to examine what can and cannot be inferred from endpoint assays, rather than as a fully biophysical description of osteogenic culture. The model is two-dimensional and does not include diffusion, matrix turnover, or mechanical feedback, all of which can influence osteogenesis in vitro [33, 50]. In this culture setting, however, diffusion is likely of secondary importance, as ascorbic acid is unstable in medium (half-life ~8 h) [34], and measured outputs primarily reflect cell-associated activity and locally retained matrix [51]. Dex and MK-4 were represented as phenomenological modulators of osteogenic activity, although both may also affect proliferation and lineage commitment through mechanisms not included here [36, 37]. Fitted parameters should therefore be interpreted as effective quantities rather than uniquely identifiable biological constants. A more detailed model would require substantially richer experimental data for calibration; without such con-straints, added realism would mainly introduce unidentifiable parameters [52], consistent with the inability to capture inhibitory responses at high dexamethasone concentrations.

Relative to earlier osteogenesis models, OsteoMin explicitly treats experimental assays as partial observations of a stochastic cell-state process. Previous cellular automaton models reproduced spatial mineralization patterns without systematically linking intermediate markers to assay-level observables [19], while continuum and ODE approaches capture aspects of matrix maturation but typically assume homogeneous populations [53].

These results indicate that most identifiable information lies in marker dynamics rather than endpoint values, which compress the underlying progression and limit mechanistic resolution. Viewed as a partially observed inverse problem, endpoint assays do not identify the underlying dynamics but instead define a set of admissible trajectories. The role of the model is therefore not to recover a unique mechanism, but to map this space of consistent dynamics and test how different hypotheses or perturbations project onto observable endpoints. Within this framework, separability depends on when and what is measured, indicating that experimental design—particularly assay timing—can be optimized to better resolve otherwise indistinguishable mechanisms.

## 6 Conclusion

We developed OsteoMin, an interpretable cellular automaton that structurally links stochastic osteogenic cell-state progression to commonly used endpoint assays of ALP activity, collagen deposition, and mineralization. By directly connecting latent differentiation dynamics to assay-level observables, the framework enables mechanistic interpretation of population-averaged readouts within a spatially resolved model of lineage progression.

Model results indicate that similar endpoint measurements can arise from distinct underlying trajectories, showing that standard assay panels constrain broad progression trends but often do not uniquely identify differentiation mechanisms. OsteoMin therefore provides both an interpretable model of osteogenic differentiation and a quantitative basis for assessing practical identifiability limits in endpoint-driven experiments, supporting the design of measurements that better resolve latent cell-state dynamics.

## Supporting information

S1 Text. Supplementary methods, derivations, and additional analyses

## Funding and acknowledgments

This work was funded by the EU Horizon 2020 programme (Marie Skłodowska-Curie grant No. 945371). A.A.D. acknowledges UiO:Life Science (University of Oslo) support for a research visit at the MERLN Institute, Maastricht University. A.C. acknowledges support from the Gravitation Program “Materials Driven Regeneration”, funded by the Netherlands Organization for Scientific Research (024.003.013).

## Author Contributions

AAD: conceptualization, methodology, software, investigation, formal analysis, visualization, writing – original draft. TC: methodology, software, investigation, formal analysis, visualization, writing – review & editing. CAH: methodology, investigation, formal analysis. HT: conceptualization, methodology, investigation, supervision, writing – review & editing. AC: methodology, supervision, funding acquisition, writing – review & editing. DKD: conceptualization, supervision, funding acquisition, writing – review & editing.

## Data availability

All data and source code are publicly available. Experimental data are archived in Dataverse (DOI:10.18710/FWDTNY). The OsteoMin source code and analysis scripts are available on GitHub (github.com/aliaslandemir/OsteoMin).

## Supporting information

S1 Text. Supplementary methods, derivations, parameterization, additional analyses, and supplementary figures supporting the OsteoMin model.

