## Supplementary material for "A cellular automaton model of osteogenic differentiation reveals identifiability limits of endpoint assays": S1 Text. Supplementary methods, derivations, and additional analyses

### Supplementary Information

**5 Supplementary Figures** **16**

**6 Reagents Information** **19**

### 1 Full model specification

This document provides detailed mathematical formulations, parameter estimation procedures, sensitivity analyses, and supplementary experimental figures supporting the OsteoMin model.

#### 1.1 Spatial and temporal representation

The model is implemented on a two-dimensional square lattice of size $n \times n$ representing the culture surface. Each lattice site corresponds to a spatial area of

$$\Delta x = 30\,\mu\text{m},$$

approximately the diameter of an MC3T3-E1 cell consistent with microscopy observations in MC3T3 cultures [1].

#### 1.2 Migration rule and calibration

Pre-OB migration was modeled as a stochastic random walk on a two-dimensional lattice. At each timestep ($\Delta t = 1\,\text{h}$), a cell attempts to move to a randomly selected empty Moore-neighbor site with probability $p_m$.

To reduce lattice anisotropy, the four axial directions were assigned a total probability weight of 2/3 and the four diagonal directions a total weight of 1/3. Thus, the probability of selecting a specific axial direction is $p_m/6$, while that of selecting a specific diagonal direction is $p_m/12$. For lattice spacing $a$, axial moves have squared length $a^2$ and diagonal moves have squared length $2a^2$, giving an expected squared displacement per timestep

$$\langle \Delta r^2 \rangle = 4\left(\frac{p_m}{6}\right)a^2 + 4\left(\frac{p_m}{12}\right)(2a^2) = \frac{4}{3}p_m a^2.$$

Since $\Delta t = 1\,\text{h}$, the model predicts

$$\langle r^2(t) \rangle = \frac{4}{3}p_m a^2\, t.$$

The migration probability $p_m$ was calibrated to reproduce the migration of MC3T3-E1 cells on flat substrates reported in [1]. In Fig. 5b of that study, the flat condition reaches an RMS displacement of approximately $70\,\mu$m after $\sim 16$ h. This corresponds to a mean-squared displacement of approximately

$$\langle r^2 \rangle \approx (70\,\mu\text{m})^2 = 4900\,\mu\text{m}^2.$$

Assuming approximately diffusive scaling at long times gives an effective displacement rate

$$\frac{\langle r^2 \rangle}{t} \approx \frac{4900}{16} \approx 320\,\mu\text{m}^2/\text{h}.$$

Matching this experimental scaling to the lattice model yields

$$\frac{4}{3} p_m a^2 = 320\,\mu\text{m}^2/\text{h}.$$

Using the simulation lattice spacing $a = 30\,\mu\text{m}$ gives

$$p_m = \frac{320}{(4/3) \times 30^2} \approx 0.27.$$

This value was used in all simulations.

##### 1.3 Osteogenic commitment

Pre-OB differentiation was implemented as a delayed stochastic transition. Osteogenic induction in the simulations was initiated at $t_{\text{osteo}} = 3$ days (72 h), corresponding to the experimental time point at which osteogenic conditions were applied.

Cells remain in the pre-OB state for a maturation interval $t_d$ after induction. After this delay, commitment occurs with a constant rate $k_{\text{commit}}$. The cumulative probability that a cell has committed by time $t$ is

$$f_{OB}(t) = \begin{cases} 0, & t < t_{\text{osteo}} + t_d, \\ 1 - e^{-k_{\text{commit}}(t - t_{\text{osteo}} - t_d)}, & t \geq t_{\text{osteo}} + t_d. \end{cases}$$

The delay parameter reflects the experimentally observed lag between osteogenic induction and early differentiation markers in MC3T3-E1 cultures. Transcriptomic analysis indicates that the transition from proliferation to osteogenic programs occurs around day 2–4 following induction [2]. We therefore set the maturation delay to $t_d = 3$ days (72 h), consistent with our experimental protocol in which osteogenic supplementation is introduced at day 3 after seeding.

Following this delay, commitment occurs with constant rate $k_{\text{commit}} = 0.0125\,\text{h}^{-1}$, corresponding to $0.30\,\text{day}^{-1}$. This implies a characteristic commitment timescale of approximately

10 days after the maturation delay:

$$T_{95} = \frac{-\ln(0.05)}{k_{\text{commit}}} \approx 10 \text{ days}.$$

Thus, most cells transition to the OB state approximately 10 days after the delay period, consistent with reported timing of MC3T3-E1 differentiation and mineralization onset between days 14–21 following osteogenic induction [2, 3].

The resulting temporal evolution of pre-OB and OB population counts under baseline osteogenic conditions is shown in Fig. 1.

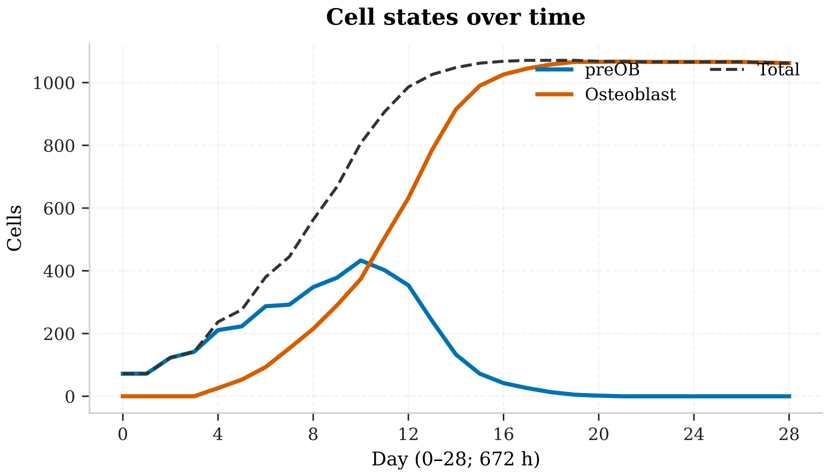

Figure 1: Temporal progression of pre-OB and OB population counts in the spatial OsteoMin model under baseline osteogenic conditions.

#### 1.4 Ascorbic acid dynamics

Ascorbic acid (AA) is supplied with osteogenic medium exchange and undergoes first-order oxidative degradation between exchanges.

The decay constant is

$$k_{AA} = \frac{\ln 2}{t_{1/2}^{AA}}, \qquad t_{1/2}^{AA} \approx 8 \text{ h}.$$

Medium is replaced every $m$ days, corresponding to an exchange interval

$$\tau_m = 24m \text{ h}.$$

AA supplementation begins simultaneously with osteogenic induction at $t_{\text{osteo}}$.

Define the medium-refresh indicator

$$u_{\text{med}}(t_k) = \begin{cases} 1, & u_{\text{osteo}}(t_k) = 1 \text{ and } \operatorname{mod}(t_k - t_{\text{osteo}}, \tau_m) = 0, \\ 0, & \text{otherwise.} \end{cases}$$

Between medium exchanges, AA decays exponentially:

$$c_{AA}(t) = C_{AA,0} \exp(-k_{AA}(t - t_p)), \qquad t \geq t_p, \tag{1}$$

where $t_p$ denotes the most recent medium exchange time.

The discrete-time update consistent with timestep $\Delta t = 1$ h is

$$c_{AA}(t_{k+1}) = c_{AA}(t_k) e^{-k_{AA}\Delta t} + C_{AA,0}\, u_{\text{med}}(t_k). \tag{2}$$

Initial condition:

$$c_{AA}(t_k) = 0, \qquad t_k < t_{\text{osteo}}.$$

Here $C_{AA,0}$ denotes the AA concentration supplied at each medium exchange. The resulting trajectory represents repeated AA pulses followed by exponential decay across successive exchange cycles. The corresponding time course of effective ascorbic acid availability under repeated medium exchange is shown in Fig. 2.

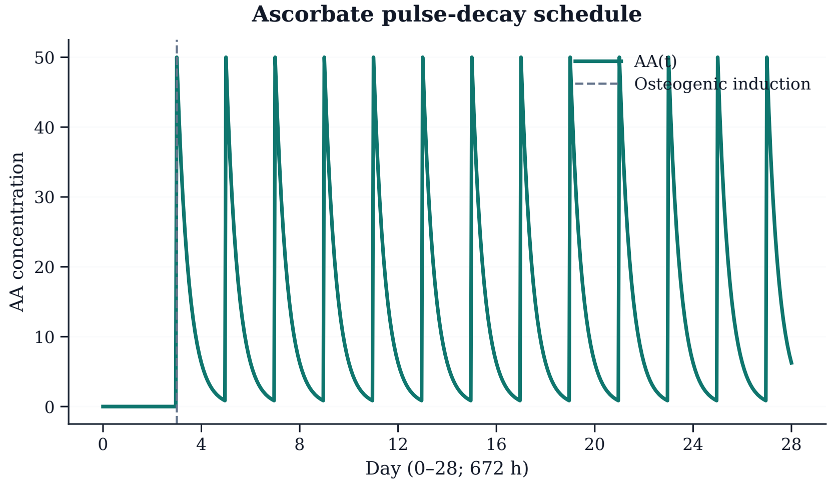

Figure 2: Effective ascorbic acid concentration in the medium under repeated supplementation and exponential degradation.

### 2 Parameterization and calibration

Published time-course and dose–response datasets were used to estimate effective phenomenological parameters describing marker production and compound modulation. Functional forms were selected to reproduce dominant experimental trends while limiting the number of poorly identifiable parameters.

#### 2.1 Basal and osteogenic marker kinetics

Early osteogenic markers such as alkaline phosphatase (ALP) activity and collagen deposition exhibit approximately monotonic increases over the experimentally observed time windows. These dynamics were therefore approximated using constant-rate accumulation models:

$$M(t) = M_0 + rt \tag{3}$$

where $M(t)$ denotes marker signal, $M_0$ the baseline level, and $r$ an effective production rate. Separate basal ($r_b$) and osteogenic ($r_o$) contributions were estimated from literature datasets.

ALP production rates were estimated from measurements at days 3, 7, 14, and 21 reported in [4]. Least-squares fitting yielded

$$r_{b,A} = 9.49 \times 10^{-2}\,\mathrm{AU\,h^{-1}}, \qquad r_{o,A} = 8.10 \times 10^{-1}\,\mathrm{AU\,h^{-1}},$$

with fit errors RMSE = 5.52 AU (basal) and 32.1 AU (osteogenic), indicating that the linear approximation captures the overall temporal increase in ALP activity over the observed interval.

Collagen production rates were estimated from measurements obtained under basal and ascorbic-acid-supplemented conditions at days 1–12 [5]. Least-squares fitting yielded

$$r_{b,C} = 3.36 \times 10^{-3}\,\mathrm{AU\,h^{-1}}, \qquad r_{o,C} = 1.30 \times 10^{-2}\,\mathrm{AU\,h^{-1}},$$

with RMSE = $9.14 \times 10^{-2}$ AU (basal) and $3.79 \times 10^{-1}$ AU (osteogenic). The approximately linear increase in collagen signal across the measured time window supports the constant-rate approximation.

#### 2.2 Mineralization dynamics

Mineral accumulation exhibits delayed onset followed by nonlinear increase reflecting extracellular matrix maturation and hydroxyapatite nucleation. Alizarin Red S (ARS) time-course measurements reported in [6] were approximated using a Hill-type activation model:

$$H(t) = H_0 + r_{o,H} \frac{t^{n_H}}{\tau_H^{n_H} + t^{n_H}} \tag{4}$$

where $\tau_H$ defines the characteristic activation timescale and $n_H$ controls transition steepness.

Fitting to measurements at days 4, 7, 10, 14, 21, and 28 yielded

$$r_{o,H} = 1.03 \times 10^{-1}\,\mathrm{AU\,h^{-1}}, \qquad \tau_H = 384\,\mathrm{h}, \qquad n_H = 2,$$

with RMSE = 2.25 AU. The Hill formulation reproduces the delayed onset and subsequent nonlinear increase in ARS signal commonly observed during MC3T3-E1 differentiation.

#### 2.3 Dex and MK-4 compound–response relationships

Dose–response effects of dexamethasone (Dex) and vitamin $K_2$ (MK-4) were estimated from published single-agent datasets using nonlinear least-squares regression. Experimental studies frequently report biphasic responses to osteogenic supplements, with intermediate concentrations producing maximal marker expression.

Dose-dependent modulation was therefore approximated using a log-parabolic response function applied multiplicatively to baseline marker production:

$$y(c) = y_0 \left[ 1 + a \left( 1 - b \left( \log_{10} \frac{c}{c_0} \right)^2 \right) \right] \tag{5}$$

where $c$ denotes compound concentration, $c_0$ the concentration associated with maximal response, $a$ the response amplitude, and $b$ controls response width in log-dose space.

Dex-dependent ALP activity was fitted to measurements at 0–100 nM [7], yielding

$$c_{D,0,A} = 22.76\,\mathrm{nM}, \quad a_{D,A} = 3.72, \quad b_{D,A} = 1.75.$$

Dex-dependent collagen response was fitted to measurements at 0–10 nM [8], yielding

$$c_{D,0,C} = 17.27\,\mathrm{nM}, \quad a_{D,C} = 2.65, \quad b_{D,C} = 0.814.$$

Dex-dependent mineralization was fitted to measurements at 0–100 nM [9], yielding

$$c_{D,0,H} = 67.79\,\text{nM}, \quad a_{D,H} = 1.15, \quad b_{D,H} = 0.345.$$

MK-4-dependent ALP activity was fitted to measurements at 0–10 $\mu$M [10], yielding

$$c_{K,0,A} = 0.7829\,\mu\text{M}, \quad a_{K,A} = 1.095, \quad b_{K,A} = 1.058.$$

Quantitative MK-4 collagen dose–response data were available only for control and 10 $\mu$M [11]. Because these measurements do not independently constrain response curvature, shape parameters were fixed to mineralization-derived values ($a_{K,C} = 4.663$, $b_{K,C} = 2.145$), yielding

$$c_{K,0,C} = 0.3480\,\mu\text{M}.$$

MK-4-dependent mineralization was fitted to measurements at 0–10 $\mu$M [10], yielding

$$c_{K,0,H} = 0.6692\,\mu\text{M}, \quad a_{K,H} = 4.665, \quad b_{K,H} = 2.150.$$

For sparse dose–response datasets ($n \leq p$), fitted coefficients should be interpreted as effective phenomenological calibration parameters rather than uniquely identifiable mechanistic constants.

Table 1: **Literature-derived and fitted production/modulation parameters.**

| **Fit** | $n$ | DoF | RMSE | **Estimated parameters** | **Source** |
|---|---|---|---|---|---|
| Dex–ALP prod. | 4 | 0 | $5.43 \times 10^{-12}$ | $c_{D,0,A} = 22.76\,\text{nM},\ a_{D,A} = 3.72,\ b_{D,A} = 1.75$ | [7] |
| Dex–coll. prod. | 3 | $NA$ | $3.02 \times 10^{-9}$ | $c_{D,0,C} = 17.27\,\text{nM},\ a_{D,C} = 2.65,\ b_{D,C} = 0.814$ | [8] |
| Dex–mineral dep. | 3 | $NA$ | $5.13 \times 10^{-9}$ | $c_{D,0,H} = 67.79\,\text{nM},\ a_{D,H} = 1.15,\ b_{D,H} = 0.345$ | [9] |
| MK-4–ALP prod. | 4 | 0 | $2.04 \times 10^{-15}$ | $c_{K,0,A} = 0.7829\,\mu\text{M},\ a_{K,A} = 1.095,\ b_{K,A} = 1.058$ | [10] |
| MK-4–coll. prod. | 2 | 0 | $7.85 \times 10^{-15}$ | $c_{K,0,C} = 0.3480\,\mu\text{M},\ a_{K,C} = 4.663,\ b_{K,C} = 2.145$ | [11] |
| MK-4–mineral dep. | 4 | 0 | $2.84 \times 10^{-14}$ | $c_{K,0,H} = 0.6692\,\mu\text{M},\ a_{K,H} = 4.665,\ b_{K,H} = 2.150$ | [10] |

Compound–response parameters were weakly constrained because only limited literature measurements were available and should therefore be interpreted as effective phenomenological coefficients rather than uniquely estimated biological constants.

#### 2.4 General parameter choices

Ascorbic acid (AA) availability was modeled as an exponentially decaying signal reflecting its rapid degradation in cell culture medium. The initial concentration was set to $50\,\mu$g/mL corresponds to 0.284 mM AA, consistent with standard osteogenic culture conditions, and decayed with an effective half-life of 8 h, consistent with reported ascorbate instability in culture media [12]. Collagen production was modulated by AA availability using a saturating gate with half-saturation constant $K_{A,C} = 5\,\mu\text{g/mL}$.

Delayed collagen support for mineralization is controlled by the maturation timescale $\tau_{\text{col}}$. Experimental studies show that extracellular matrix deposition precedes mineral formation and that hydroxyapatite nucleation requires a sufficiently developed collagen scaffold [5, 13]. We therefore set $\tau_{\text{col}} = 48$ h as an effective matrix maturation timescale.

Because collagen provides nucleation sites for hydroxyapatite formation, mineral deposition depends on extracellular matrix availability [13]. This dependence is represented phenomenologically using a saturating function with half-saturation constant $K_{\text{col}} = 1.0$ in normalized collagen units, defining the matrix level at which mineral deposition reaches half-maximal rate.

The parameters $\tau_H$ and $n_H$ determine the characteristic timescale and steepness of mineralization activation following OB commitment. To reproduce the delayed onset and progressive increase in Alizarin Red S staining observed during MC3T3-E1 differentiation, we fixed $\tau_H = 384\,\text{h}$ and $n_H = 2$. The osteogenic mineral deposition rate was set to $r_{o,H} = 0.103\,\text{AU}\,\text{h}^{-1}$, obtained from fitting mineral accumulation dynamics reported in [6].

#### 2.5 Seeding estimates

Cell concentration was determined experimentally by hemocytometer counting following a 10-fold dilution. A total of 49 cells were counted across 12 large squares, corresponding to an average of $49/12 \approx 4.1$ cells per square. Since each large square represents $10^{-4}\,\text{mL}$, the diluted suspension concentration was $4.1 \times 10^4$ cells/mL. After correcting for dilution, the original cell concentration was estimated as $4.1 \times 10^5$ cells/mL.

Each well received $20\,\mu\text{L}$ of the cell suspension, corresponding to approximately $8.0 \times 10^3$ cells per well.

Cells were seeded onto 12 mm diameter glass coverslips with a surface area $A = \pi r^2 \approx 1.13\,\text{cm}^2$. This yields an initial seeding density of

$$\rho \approx 7.1 \times 10^3 \text{ cells/cm}^2 \quad (\approx 72 \text{ cells/mm}^2).$$

#### 3 Sensitivity and perturbation analyses

All parameters were sampled independently from uniform distributions using Saltelli sampling.

Table 2: Parameter ranges used for Sobol sensitivity analysis.

| Parameter | Description | Lower | Upper |
|---|---|---|---|
| ALP basal prod. | ALP basal production rate | 0.07 | 0.13 |
| ALP osteogenic prod. | ALP osteogenic production rate | 0.60 | 1.05 |
| Collagen basal prod. | Collagen basal production rate | 0.0020 | 0.0055 |
| Collagen osteogenic prod. | Collagen osteogenic production rate | 0.0080 | 0.0200 |
| Mineral osteogenic dep. | Mineral osteogenic deposition rate | 0.075 | 0.150 |
| Mineral Hill n | Mineral Hill function $n$ | 1.5 | 3.5 |
| Mineralization tau | Mineralization activation timescale | 288.0 | 480.0 |
| Collagen maturation tau | Collagen maturation timescale ($\tau_{\mathrm{col}}$) | 24.0 | 72.0 |
| Collagen support K | Collagen support half-saturation ($K_{\mathrm{col}}$) | 0.5 | 2.0 |
| Commitment rate | Commitment rate | 0.0075 | 0.0200 |
| Pre-OB apoptosis prob. | Pre-OB apoptosis probability | 0.0005 | 0.0015 |
| Cell motility prob. | Cell motility probability | 0.15 | 0.40 |
| Ascorbate half-life | Ascorbate half-life | 4.0 | 12.0 |
| Collagen AA K | AA half-saturation for collagen ($K_{A,C}$) | 2.5 | 8.0 |
| Commitment delay | Commitment delay | 24.0 | 72.0 |
| Min division interval | Minimum division interval | 30.0 | 46.0 |
| Division period | Division period | 8.0 | 24.0 |
| Division prob. | Division attempt probability | 0.75 | 1.0 |
| Osteogenic induction | Osteogenic induction time | 48.0 | 96.0 |
| Media refresh interval | Media refresh interval | 24.0 | 72.0 |

##### 3.1 Structural ablation and nonspatial comparator

We assessed the contribution of key mechanisms in the spatial osteogenesis model by systematically removing individual processes and comparing results with a matched nonspatial (well-mixed) formulation. Analyses correspond to condition C2 (Dex = 0, MK-4 = 0).

The reference system was the full spatial model initialized from the baseline configuration. Simulations were performed for 28 days using 10 replicate seeds. Endpoints were ALP (day 10), collagen (day 14), mineralization (day 28), and total cell number (day 28).

###### 3.1.1 Nonspatial comparator

To isolate the contribution of spatial organization, we constructed a nonspatial population-level formulation in which the explicit lattice representation of the agent-based model (ABM) is removed while preserving the same kinetic parameterization and assay mappings. Initial conditions were matched by setting the starting pre-OB and OB population counts equal to the day-0 reference state of the spatial simulation. The effective population capacity was fixed to $K = 1089$, corresponding to the number of lattice sites in the spatial model ($33 \times 33$ grid representing $1\,\text{mm} \times 1\,\text{mm}$). Model outputs were evaluated at the same assay-aligned time points as the spatial simulations (day 10 ALP, day 14 collagen, day 28 mineral, and day-28 total cell number). Because the nonspatial formulation does not include stochastic cell-level events, replicate variability was introduced by applying log-normal perturbations (coefficient of variation 0.10) to selected kinetic rate parameters.

In the nonspatial formulation, cells are not assigned spatial positions. Instead, the numbers of pre-OB and OB cells are represented as total population counts that evolve in time according to the same kinetic rate parameters used in the spatial model. Cell proliferation is limited by total population size relative to the carrying capacity $K$, replacing the spatial occupancy constraint that restricts daughter-cell placement in the ABM lattice. Differentiation commitment from pre-OB to OB is implemented as a rate-dependent population transfer after the same induction delay used in the spatial model, but without stochastic per-cell transitions. As in the spatial model, cell division contributes newly generated cells to the pre-OB population regardless of parent state, preserving the same lineage structure.

Extracellular matrix production and maturation follow identical kinetic expressions as in the spatial model but are evaluated using population-averaged quantities rather than spatially resolved fields. Consequently, mineral formation is determined by the global availability of mature collagen and the average progression of OB cells beyond commitment, rather than requiring spatial co-localization of osteoblast occupancy and sufficiently matured matrix at individual lattice sites. This formulation preserves the same biochemical rate structure and assay observables as the spatial ABM while removing spatial constraints on cell placement, local crowding effects, and stochastic variation in commitment timing. Differences between the two formulations therefore reflect the contribution of spatial exclusion, spatially heterogeneous matrix accumulation, and asynchronous single-cell state transitions to emergent mineralization dynamics.

##### 3.2 Selected Dex/MK-4 dose ranges for analysis

Dose selections were defined for single-compound variations and combined Dex–MK-4 response mapping.

Table 3: **Single-compound dose ranges.** One compound was varied while the other was fixed at zero. The baseline condition corresponds to Dex = 0 nM and MK-4 = 0 $\mu$M.

| Condition | Dose values |
|---|---|
| MK-4 only | Dex = 0 nM; 0.03, 0.10, 0.20, 0.30, 0.60, 1.00, 2.50, 5.00, 10 $\mu$M |
| Dex only | MK-4 = 0 $\mu$M; 0.3, 1, 2, 3, 6, 10, 20, 30, 60, 100 nM |
| Baseline | Dex = 0 nM; MK-4 = 0 $\mu$M |

Table 4: **Combined Dex–MK-4 dose ranges.** All pairwise combinations of the listed Dex and MK-4 concentrations were evaluated, yielding 100 total conditions.

| Compound | Dose values |
|---|---|
| Dex | 0, 1, 5, 10, 20, 30, 40, 60, 80, 100 nM |
| MK-4 | 0, 0.1, 0.3, 0.5, 1, 2, 3, 5, 7, 10 $\mu$M |

##### 3.3 Perturbation-family identifiability analysis

To assess whether assay-aligned endpoints distinguish mechanistic perturbations, parameters were varied within biologically plausible ranges using family-wise quasi-random sampling. For each perturbation family, only the associated parameters were varied, while all others were fixed at baseline values. Sampling schemes were adapted to parameter type and structural constraints.

#### 4 Spatial and image-based analyses

##### 4.1 Spatial autocorrelation

Let $H(x, y)$ denote the final mineral field on our grid. The mean-subtracted field is

$$\tilde{H}(x, y) = H(x, y) - \bar{H}, \qquad \bar{H} = \frac{1}{WH} \sum_{x,y} H(x, y). \tag{6}$$

The unnormalized autocovariance was computed using the Wiener–Khinchin theorem

$$C_{\text{raw}}(\Delta x, \Delta y) = \text{IFFT2}\left( \left| \text{FFT2}(\tilde{H}) \right|^2 \right), \tag{7}$$

and normalized as

$$C(\Delta x, \Delta y) = \frac{C_{\text{raw}}(\Delta x, \Delta y)}{C_{\text{raw}}(0, 0)}. \tag{8}$$

Table 5: Parameter ranges used for perturbation-family identifiability analysis. Each family was varied independently around the baseline parameter set.

| Parameter | Description | Lower | Upper |
|---|---|---|---|
| **Spatial pattern** | | | |
| init_slug | Initial spatial seeding motif | C01 | C50 |
| boundary_scale | Spatial scaling factor | fixed | fixed |
| **Cell number** | | | |
| init_cells | Initial number of pre-OB cells | 27 | 212 |
| grid_spacing | Structured seeding grid spacing | 1 | 4 |
| **Cell dynamics** | | | |
| osteo_induction_hours | Osteogenic induction time (h) | 50 | 95 |
| commit_delay | Commitment delay $t_d$ (h) | 24 | 71 |
| commit_k | Commitment rate $k_{\text{commit}}$ | 0.00825 | 0.01970 |
| move_prob | Cell motility probability $p_m$ | 0.1536 | 0.3929 |
| min_div_hours | Minimum division interval (h) | 32 | 45 |
| max_div_hours | Maximum division interval (h) | 43 | 62 |
| apoptosis_prob | Pre-OB apoptosis probability | 0.000547 | 0.001451 |
| **Ascorbic acid / media chemistry** | | | |
| aa0 | Initial AA concentration $C_{AA,0}$ (µg/mL) | 25.72 | 89.13 |
| aa_half_life | Ascorbate half-life $t_{1/2}^{AA}$ (h) | 4.07 | 11.94 |
| collagen_aa_km | AA half-saturation for collagen $K_{A,C}$ | 2.58 | 7.92 |
| media_interval | Media refresh interval $\tau_m$ (h) | 24 | 72 |

Radial averaging was performed using periodic distances

$$d_x = \min(\Delta x,\, W - \Delta x), \qquad d_y = \min(\Delta y,\, H - \Delta y), \tag{9}$$

$$r = \sqrt{d_x^2 + d_y^2}, \tag{10}$$

with integer radial bins $i \in [0, \lfloor \min(W, H)/2 \rfloor)$. The radial correlation profile $C_r(i)$ was obtained by averaging $C(\Delta x, \Delta y)$ within each annulus.

The autocorrelation length $\xi$ was defined as the first radius where $C_r(i)$ crossed $e^{-1}$, computed by linear interpolation between adjacent bins:

$$C_r(i-1) \geq e^{-1}, \qquad C_r(i) \leq e^{-1}. \tag{11}$$

For each condition, the reported value corresponds to the replicate mean

$$\xi = \mu_\xi \pm \sigma_\xi, \qquad \mu_\xi = \frac{1}{n}\sum_{m=1}^{n} \xi_m. \tag{12}$$

Distances are reported in lattice units.

Spatial heterogeneity of mineral deposition quantified from ARS images is summarized in Fig. 3.

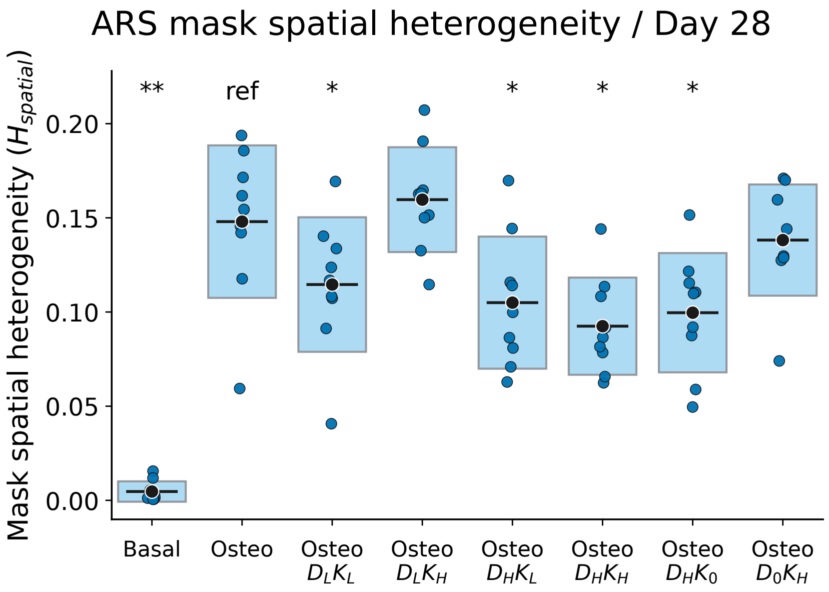

Figure 3: Spatial heterogeneity of mineral deposition at day 28 quantified from ARS images.

Spatial organization was characterized using a sliding-window local variance metric ($64 \times 64$

px), where heterogeneity ($H_{\text{spatial}}$) was defined as the image-averaged local variance. Replicate values ($n = 9$ per condition) are reported as mean $\pm$ SD. Statistical differences relative to the osteogenic reference condition (C2) were evaluated using two-sided Mann–Whitney U tests with Benjamini–Hochberg correction following an overall Kruskal–Wallis test. Representative spatial distributions of the simulated mineral field at day 28 across conditions are shown in Fig. 4.

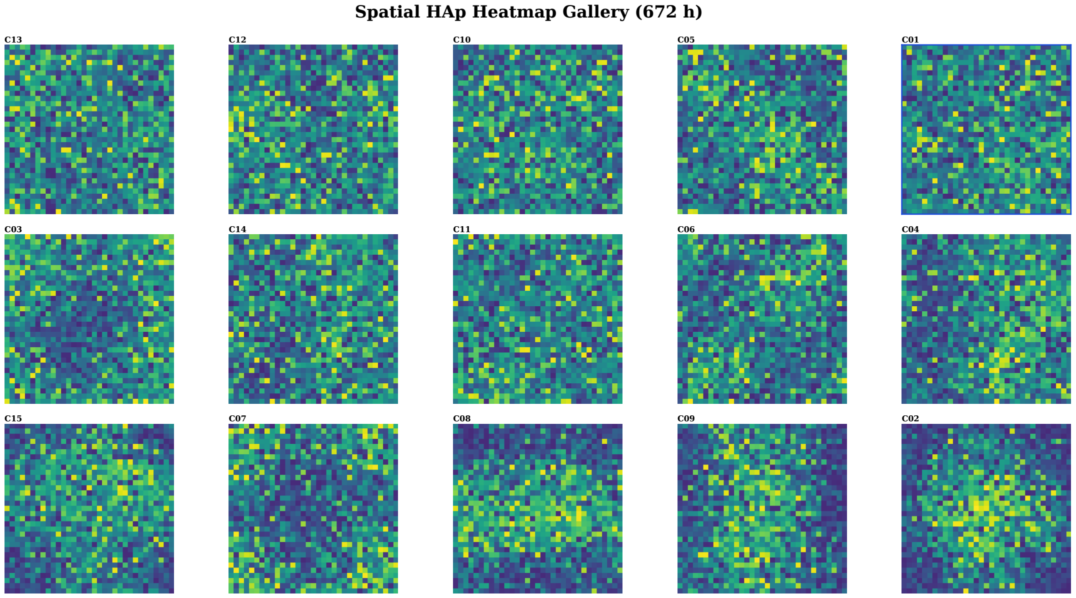

Figure 4: Spatial distributions of the simulated mineral field at day 28 across conditions.

High Dex conditions showed reduced spatial heterogeneity, indicating more spatially uniform mineral distribution, consistent with trends observed in nuclear morphology (DAPI area) from confocal measurements.

#### 4.2 Nuclear morphology analysis

DAPI-stained images were segmented by intensity thresholding to quantify nuclear projected area and aspect ratio, summarized in Figs. 5 and 6, respectively.

Nuclear morphology was analyzed from DAPI-stained images to assess condition-dependent changes in cell organization during osteogenic progression. Increases in nuclear projected area and aspect ratio indicate cytoskeletal remodeling and more elongated cell states associated with differentiation and matrix production.

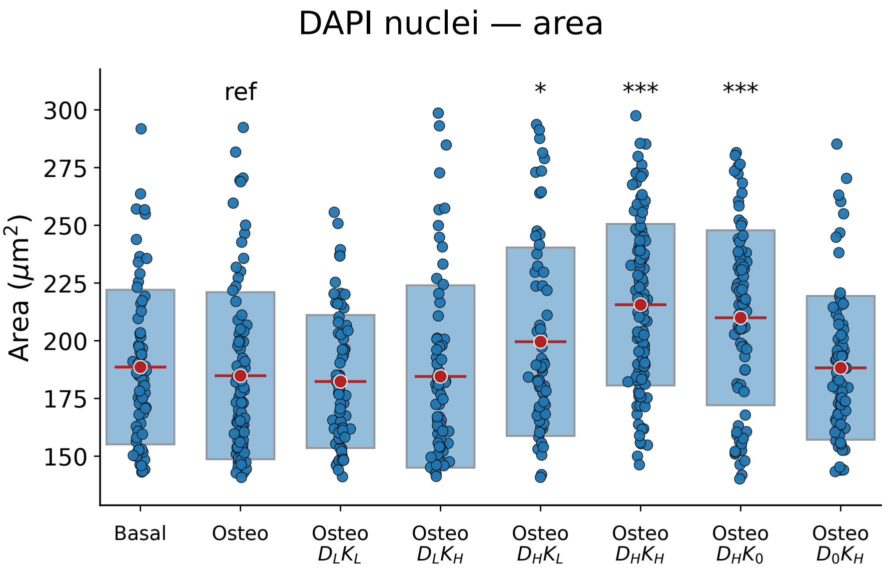

Figure 5: Nuclear projected area measured from DAPI-stained images across culture conditions.

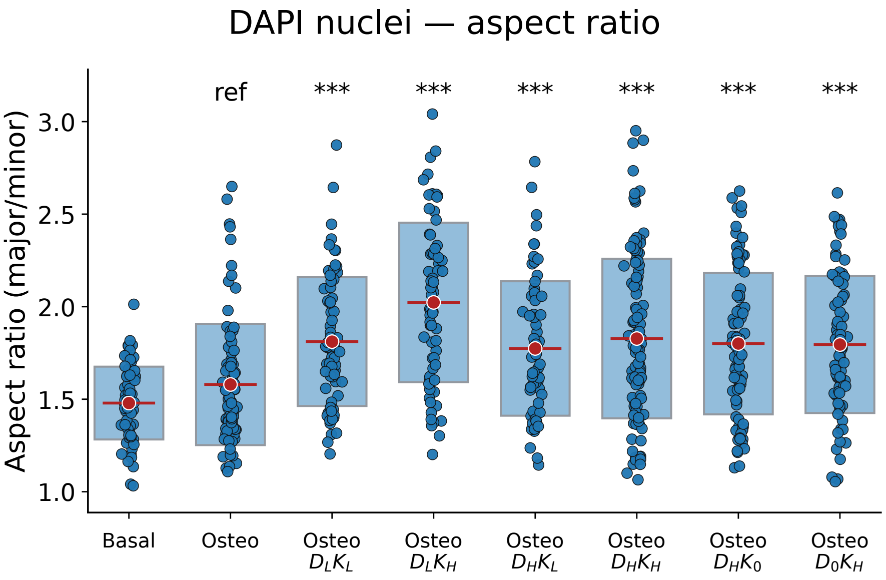

Figure 6: Nuclear aspect ratio (major/minor axis) measured from DAPI-stained images across culture conditions.

#### 5 Supplementary Figures

Representative brightfield images across all experimental conditions are shown in Fig. 7. Differences in texture and local density reflect condition-dependent changes in proliferation, morphology, and matrix formation during osteogenic progression.

Representative Alizarin Red S (ARS) images at day 28 across all experimental conditions are shown in Fig. 8. Spatial patterns indicate heterogeneous mineral deposition with localized nodular regions and condition-dependent differences in coverage and intensity.

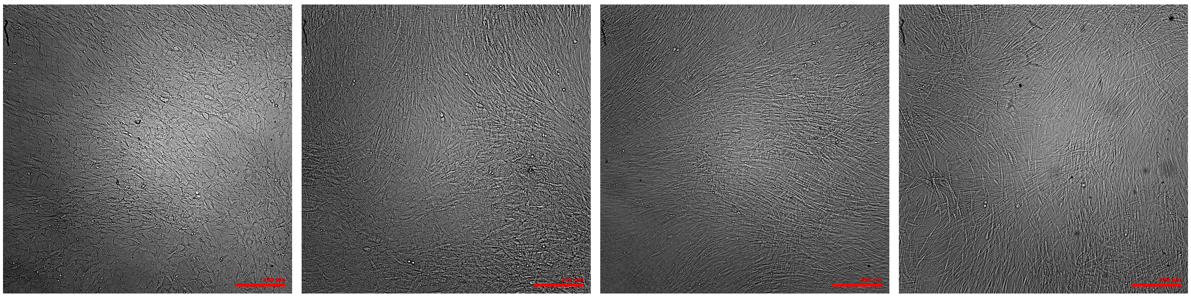

(a) Basal (b) Osteo (c) Osteo + $D_L$ $K_L$ (d) Osteo + $D_L$ $K_H$

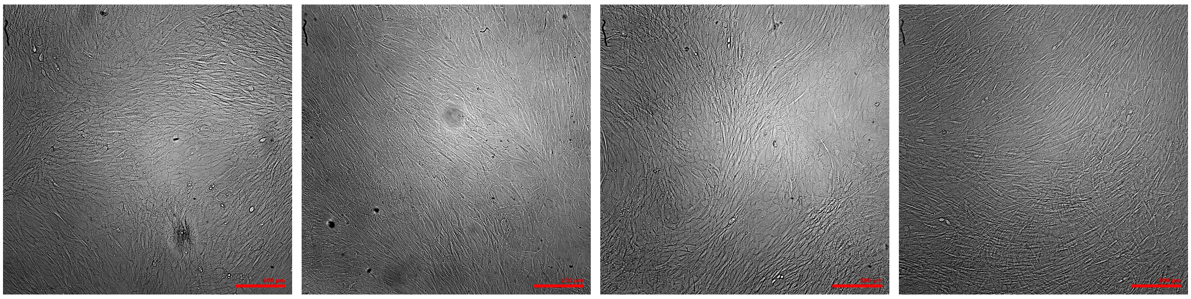

(e) Osteo + $D_H$ $K_L$ (f) Osteo + $D_H$ $K_H$ (g) Osteo + $D_H$ $K_0$ (h) Osteo + $D_0$ $K_H$

Figure 7: Representative brightfield images acquired during culture across experimental conditions C1–C8.

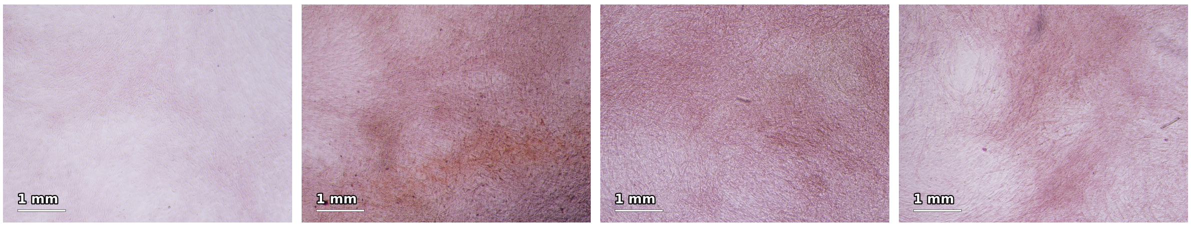

(a) Basal (b) Osteo (c) Osteo + $D_L$ $K_L$ (d) Osteo + $D_L$ $K_H$

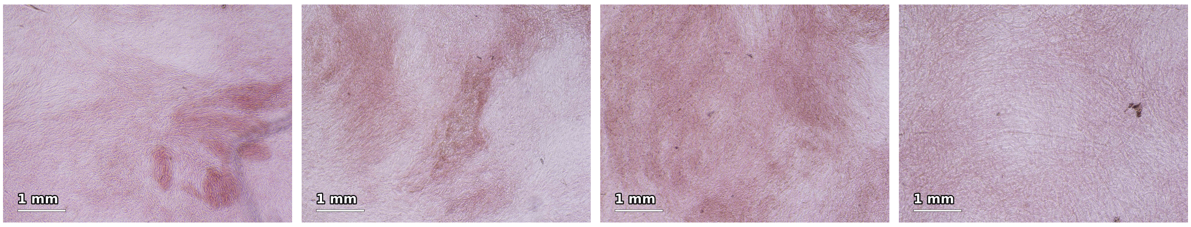

(e) Osteo + $D_H$ $K_L$ (f) Osteo + $D_H$ $K_H$ (g) Osteo + $D_H$ $K_0$ (h) Osteo + $D_0$ $K_H$

Figure 8: Representative Alizarin Red S (ARS) images at day 28 across experimental conditions C1–C8.

The sensitivity of ARS-positive area quantification to the selected threshold is shown for a representative C2 image in Fig. 9.

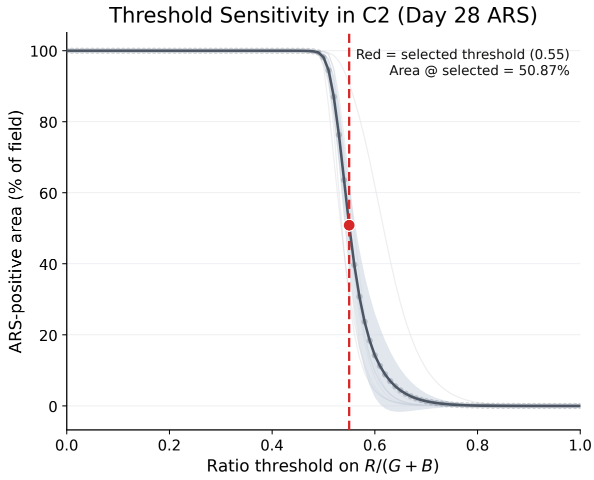

Figure 9: Threshold sensitivity analysis for ARS-positive area quantification in condition C2 at day 28. The dashed red line indicates the selected threshold (0.55), corresponding to an ARS-positive area fraction of 50.87% for the representative image. Shaded curves indicate replicate-level variation.

### 6 Reagents Information

| **Reagent** | Catalog No. | Supplier | Notes |
|---|---|---|---|
| **MC3T3-E1 SubClone 4** | CRL-2593 | ATCC | Mouse cell line. |
| **$\alpha$-MEM (no ascorbic acid)** | A10490-01 | Gibco (Thermo Fisher) | Basal medium. |
| **Fetal Bovine Serum (FBS)** | F7524-500ML | Sigma-Aldrich | 10% v/v supplement. |
| **Antibiotic–Antimycotic** | A5955-100ML | Sigma-Aldrich | 1% v/v supplement. |
| **DPBS (no Ca, Mg)** | 14190136-100ML | Thermo Fisher | Washing buffer. |
| **Triton X-100** | 11498696 | Thermo Fisher | Lysis detergent. |
| **BSA** | A9647 | Sigma-Aldrich | Blocking reagent. |
| **Absolute Ethanol (99.5%)** | 733194 | Avantor / VWR | Solvent. |
| **Mowiol 4-88** | 17951-1006 | Polysciences | Mounting medium. |
| **Paraformaldehyde (PFA)** | 28908 | Thermo Fisher | 4% fixative. |
| **TRIS Ultra Pure** | 0497-1KG | VWR / Avantor | Buffer base. |
| **$MgCl_2 \cdot 6H_2O$** | AO344733 230 | Merck / Avantor | Analytical grade. |
| **NaOH** | B1155898 517 | Merck / Avantor | pH adjustment. |
| **L-Ascorbic Acid (AA)** | A4403 | Sigma-Aldrich | Supplement. |
| **$\beta$-Glycerophosphate ($\beta$-GP)** | G9422 | Sigma-Aldrich | Supplement. |
| **Dexamethasone (Dex)** | D4902 | Sigma-Aldrich | Glucocorticoid. |
| **Vitamin $K_2$ (MK-4)** | V9378 | Sigma-Aldrich | Cofactor. |
| **Alizarin Red S (ARS)** | A5533 | Sigma-Aldrich | Mineral stain. |
| **Alkaline Phosphatase (pNPP)** | N1891 | Sigma-Aldrich | Colorimetric assay. |
| **Mouse anti-Collagen I** | MA1-26771 | Thermo Fisher | Primary antibody. |
| **Goat anti-Mouse IgG Alexa 488** | A32723 | Thermo Fisher | Secondary antibody. |
| **Phalloidin Alexa 568** | A12380 | Thermo Fisher | F-actin stain. |
| **DAPI** | D1306 | Thermo Fisher | Nuclear stain. |
